# Organ-specific fibroblast dynamics revealed via an integrated experimental-computational framework

**DOI:** 10.64898/2026.06.05.730427

**Authors:** Chloe L. Stewart, Yuan Yin, Maria-Alexa Cosma, P.R Riley, Sarah L. Waters, Ruth E. Baker

**Affiliations:** Institute of Developmental and Regenerative Medicine, University of Oxford, UK; Department of Physiology, Anatomy & Genetics, University of Oxford, UK; Mathematical Institute, University of Oxford, UK; Department of Cardiology, Medical University of Graz, Austria

## Abstract

Fibrosis is a progressive pathological process driven by dysregulated fibroblast activity and excessive extracellular matrix deposition, leading to scarring, functional decline and eventually organ failure. Although fibroblasts are key mediators of fibrotic remodelling, it remains unclear whether their behaviours are conserved across tissues or organ specific. Here, we combine experimental and computational approaches to dissect fibroblast dynamics underlying cardiac and pulmonary fibrosis. Using *in vitro* assays, we demonstrate that cardiac and lung fibroblasts differ in morphology, proliferation, and collagen matrix organisation, likely reflecting tissue-specific mechanical and biochemical demands. Lung fibroblasts display higher proliferative capacity and increased myofibroblast activation, forming condition-dependent collagen architectures, with fibres that are more aligned under proinflammatory and fibrotic conditions, but less so under anti-inflammatory conditions. In contrast, cardiac fibroblasts consistently generate diffuse, isotropic collagen matrices across all cytokine conditions. Integrating these data into a mechanistic computational model of fibroblast-collagen interactions, we show that fibroblast motility, rather than cell-cell interactions, dominates tissue dynamics, and that coupling between motility and local collagen density regulates matrix architecture. Simulations reveal that collagen-dependent motility and density-dependent collagen secretion prevent maladaptive matrix alignment, identifying potential mechanisms restraining fibrotic progression. Our findings uncover organ-specific fibroblast behaviours and highlight how integrating experimental and computational frameworks can illuminate the dynamic rules governing fibrosis and inform targeted antifibrotic strategies.

## Introduction

Fibrosis is a chronic and progressive condition characterised by dysregulated wound repair mechanisms, resulting in excessive accumulation and abnormal restructuring of the extracellular matrix (ECM) and ultimately leading to tissue scarring^1^. It is a major global health concern due to its ability to affect multiple organs, including the heart^2^, lungs^3^, liver^4^, kidneys^5^, and skin^6^, and it can potentially cause irreversible organ failure. Whilst fibrosis involves a complex and multifactorial pathophysiology, fibroblasts are recognised as central cellular mediators due to their essential role in ECM synthesis and remodelling^2,7^. The differentiation of activated fibroblasts into myofibroblasts is a key step in early tissue repair, due to the role of myofibroblasts in driving wound contraction and rapid synthesis of a provisional matrix^8,9^. Whilst transient fibroblast activation aids early wound healing, persistent activation under pathological conditions can lead to fibrosis through excessive ECM accumulation and myofibroblast persistence^10^.

A major constituent of the ECM involved in both tissue structure maintenance and fibrosis is collagen. More than 30 different types of collagens are known to-date, but collagens I and III, which are predominantly produced by fibroblasts and myofibroblasts, are the most extensively studied in the context of tissue repair and fibrosis^11–13^. During early wound healing, collagen III fibres are synthesised by both fibroblasts and myofibroblasts to form a flexible provisional matrix that supports cell infiltration. Collagen I fibres gradually replace this provisional matrix to restore tissue strength and stability^11^. Fibrosis is characterised by an accumulation of collagens I and III, with large amounts of collagen I causing tissue stiffening and compromised organ function^14–16^. In addition to dictating tissue mechanics, deposited collagens also modulate cell behaviour^17,18^ and provide contact guidance cues to direct cell migration and infiltration^19,20^.

Despite extensive research into the role of fibroblasts during fibrosis^1,21,22^, there are currently few antifibrotic therapies approved for clinical use, and those that are available often show limited efficacy across organs and disease contexts^23,24^. For example, whilst pirfenidone and nintedanib are approved for managing idiopathic pulmonary fibrosis^25^, no licensed antifibrotic treatments are currently available for cardiac, liver, or kidney fibrosis in the UK^26^. This raises fundamental questions as to whether there is a unifying mechanism of fibrosis across organs, or whether fibroblasts from different organs exhibit mechanistically distinct behaviours during the initiation and progression of fibrosis? Moreover, could these distinctions be exploited to develop more effective, tissue-specific antifibrotic therapies? We hypothesise that organotypic fibroblasts contribute to the diverse pathophysiology and clinical manifestations of fibrosis observed across organ systems^27^. Increasing evidence supports this concept, with fibroblast populations displaying substantial intra- and inter-tissue heterogeneity and plasticity^28–30^, and exhibiting distinct transcriptional programs^31–33^ and phenotypic profiles^25,34^. Investigating shared and organ-specific fibroblast features may reveal conserved mechanisms for broad antifibrotic approaches, whilst also capturing heterogeneity that could guide organ-specific therapeutic strategies^31^.

Attempts to modulate the fibrotic microenvironment to promote regeneration and halt disease progression have largely been unsuccessful, in part due to the complex interplay of molecular, cellular, and tissue-level factors that govern fibrosis onset and progression^35,36^. In this context, computational modelling offers a powerful means of gaining mechanistic insight. Models provide a rigorous framework to disentangle the various interacting biological processes, simulate tissue dynamics, test mechanistic hypotheses, and evaluate therapeutic scenarios across multiple spatial and temporal scales. Computational modelling has advanced our understanding of key biological processes involved in tissue repair and fibrosis ^37–40^, and validated models can serve as predictive tools for assessing potential anti-fibrotic therapeutic strategies^41,42^. A broad range of modelling approaches have been employed to explore wound healing and fibrotic mechanisms^43–46^. However, many existing models focus on a single organ or disease context and are often developed independently of experimental data, limiting their ability to uncover shared versus tissue-specific drivers of disease.

To investigate the mechanisms underlying organotypic fibrosis, we adopted an interdisciplinary approach that integrates experimental data with computational modelling to examine how tissue origin and microenvironmental cues shape fibroblast behaviour. Using a series of functional assays, we experimentally characterised the behavioural similarities and differences between phenotypically distinct heart and lung fibroblasts as drivers of cardiac and pulmonary fibrosis, respectively. Informed by these experimental data, we extended the framework of Yin et al.^47^ to develop a computational model that captures the key mechanisms involved in fibrosis, specifically fibroblast motility, proliferation, cell-cell interactions (i.e., population pressure and adhesion), fibroblast-mediated collagen secretion and degradation, and contact guidance. We demonstrate that this interdisciplinary experimental-computational framework enables the systematic testing of mechanistic hypotheses underlying the observed fibroblast behaviours and fibrotic outcome.

## Results

### Cardiac and lung fibroblasts exhibit differences in morphology, activation, and collagen deposition

We first employed a suite of *in vitro* assays to characterise baseline differences in morphology, activation state, and collagen matrix production between cardiac and lung fibroblasts. In an image-based cell morphology assay, cardiac and lung fibroblasts appeared morphologically different under baseline conditions. Cardiac fibroblasts were more expansive with irregular cell peripheries, whilst lung fibroblasts exhibited smaller, more compact shapes (Figure 1a). To assess the extent to which cardiac and lung fibroblasts exhibit directional alignment, we analysed their cytoskeletal orientation using OrientationJ. The nematic order parameter (S), where S=1 indicates perfect cell alignment across the population, was used to quantify collective cell orientation. Whilst both cardiac and lung fibroblasts exhibited broadly similar distributions of orientation angles (Figure 1a), cardiac fibroblasts showed a modest but statistically nonsignificant tendency towards alignment (S = 0.46 ± 0.20) compared to lung fibroblasts (S = 0.36 ± 0.21) (Figure 1b).

**Figure 1.**
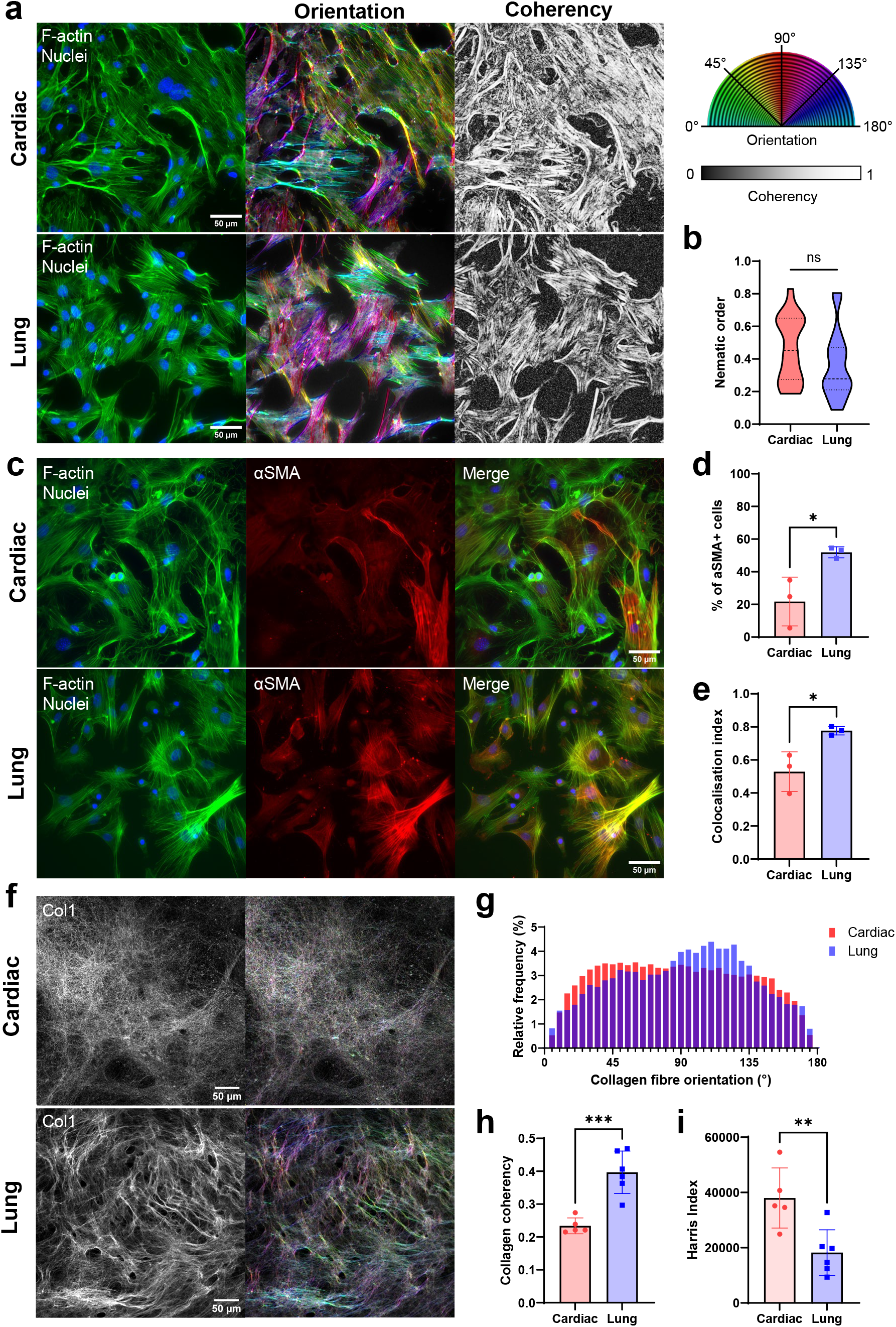
Cell morphology and expression of αSMA by cardiac and lung fibroblasts. **(a)** Cardiac and lung fibroblasts stained for F-actin filaments with spatial maps depicting F-actin filament orientation and coherency in cardiac and lung fibroblasts. The hue and saturation of the orientation colour map corresponds to F-actin orientation and coherency, respectively. **(b)** Nematic order of cardiac and lung fibroblast populations, indicating the tendency of cells to align in the same direction. Data is the distribution of 18 replicates with the median and quartiles displayed. **(c)** Expression of αSMA by fibroblasts under standard culture conditions. **(d)** Percentage of cells containing αSMA stress fibres in cardiac and lung fibroblast populations. Data is the mean of three replicates with SD. **(e)** Manders’ colocalisation index representing the proportion of F-actin filaments that overlap with αSMA stress fibres. Data is the mean of three replicates with SD. **(f)** Decellularised collagen I matrices deposited by cardiac and lung fibroblasts over 7 days and spatial maps representing fibre orientations. **(g)** Distribution of collagen fibre orientations obtained using the OrientationJ Vector Field plugin for images in (f). **(h)** Averaged collagen fibre coherency and **(i)** Harris Index for cardiac and lung-derived collagen matrices. Data is the mean of five (cardiac) and six (lung) technical replicates with SD. Statistical significance was determined by unpaired t-tests (* *p*<0.05; ** *p*<0.01; *** *p*<0.005; ns, non-significant).

To examine tissue-specific differences in fibroblast activation and differentiation to a myofibroblast state under normal culture conditions, we compared the expression of alpha-smooth muscle actin (□-SMA) stress fibres, a hallmark of myofibroblasts, across cardiac and lung fibroblasts. Lung fibroblast populations exhibited both a significantly greater proportion of □-SMA-positive cells (Figure 1c,d; p<0.05) and greater colocalisation of F-actin with □-SMA (Figure 1e; p<0.05), indicating a high alignment of □-SMA stress fibres along cytoskeletal F-actin filaments. This suggests that lung fibroblasts have a greater propensity to differentiate into myofibroblasts compared to cardiac fibroblasts.

The production of collagen I is a fundamental property of both fibroblasts and myofibroblasts, and collagen I is a major component of fibrotic tissue. Under baseline conditions, cardiac and lung fibroblasts exhibited tissue-specific differences in their collagen I matrix architecture. Cardiac fibroblasts produced a diffuse, web-like collagen network, whereas lung fibroblasts appeared to deposit collagen in thicker and more bundled fibre arrangements (Figure 1f). Collagen fibre alignment was quantified by coherency analysis using OrientationJ, whereby a coherency value of 1 indicates a perfectly uniform alignment of fibres. Lung-derived matrices exhibited significantly higher fibre alignment than cardiac-derived matrices, as indicated by a peak observed in the orientation distribution data (Figure 1g) and greater fibre coherency (Figure 1h; p<0.005).

Cardiac-derived matrices also displayed a ‘criss-cross’ architecture with frequent directional changes. Harris Corner Detection analysis was employed to distinguish points of strong pixel intensity changes due to sudden changes in fibre direction. Cardiac matrices possessed a significantly higher Harris Index, which is consistent with visual observations of increased matrix intersections and directional variance (Figure 1i; p<0.01). These structural differences between cardiac and lung-derived collagen matrices may reflect tissue-specific mechanical demands.

Collectively these experimental observations reveal fundamental differences in cardiac and lung fibroblast behaviour and collagen matrix production under normal culture conditions.

### Cardiac and lung fibroblasts exhibit similar wound closure dynamics, driven mainly by cell migration

To determine whether tissue origin influences fibroblast migration during injury and repair, we compared cardiac and lung fibroblasts using a scratch assay. Prior to scratching, lung fibroblasts had significantly larger cell areas (4430 ± 1170 µm^2^) compared to cardiac fibroblasts (2100 ± 365 µm^2^) (Figure 2a; p<0.05), which contrasted with our earlier observations (in lower density assays) that cardiac fibroblasts possessed larger cell areas than lung fibroblasts (Figure 1a). This discrepancy indicates that cardiac fibroblasts may more readily adapt their morphology to changes in the local cell density and this potentially impacts the degree to which they experience cell-cell contact during migration and proliferation.

**Figure 2.**
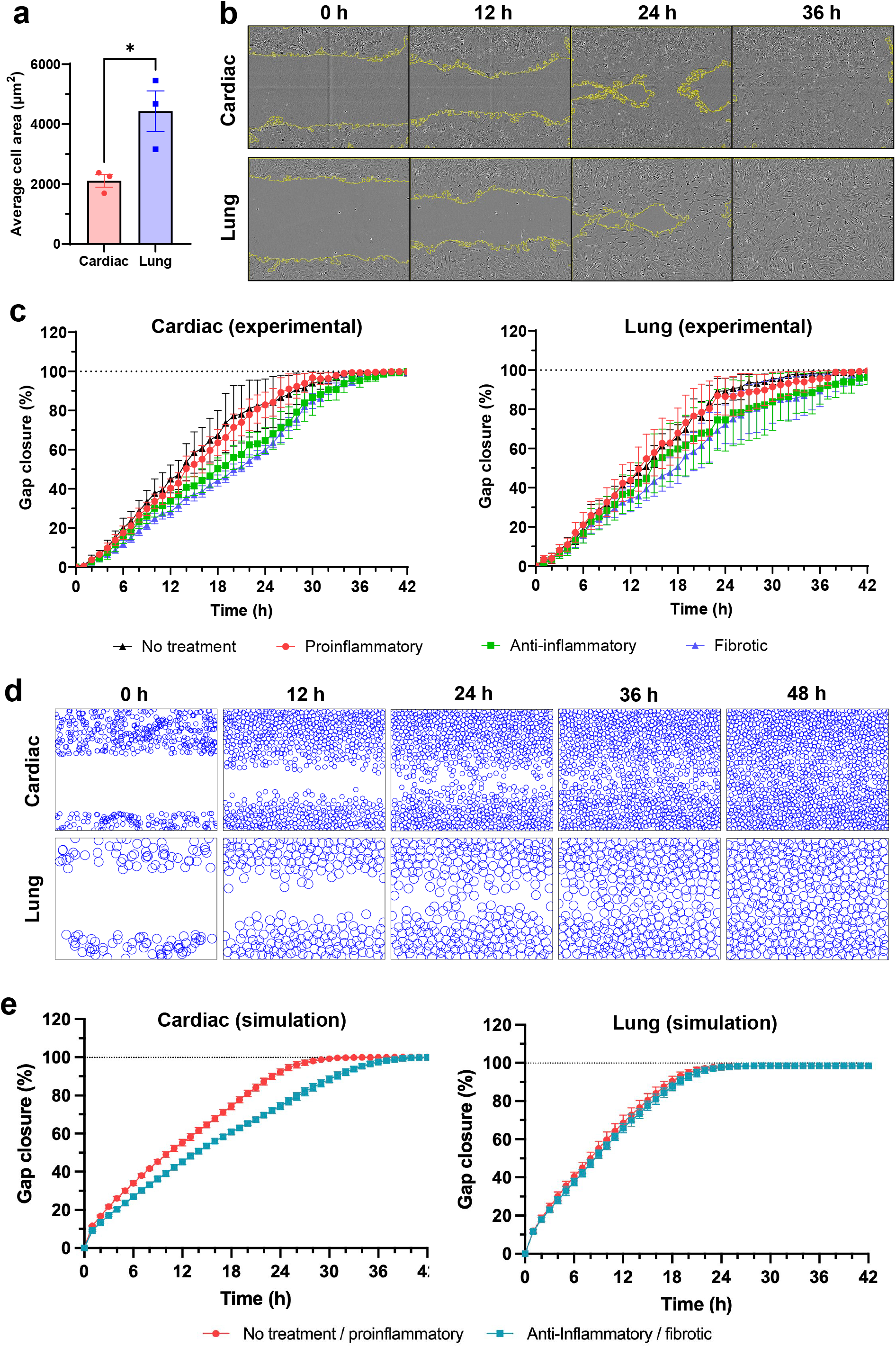
Fibroblast migration in an *in vitro* scratch assay. **(a)** Average cell size at t=0 hours. Data is the mean of three biological replicates from independent experiments with SEM. Statistical significance determined by an unpaired t-test (* *p*<0.05). **(b)** Snapshots of scratch wound closure over time by cardiac and lung fibroblasts without cytokine treatment. **(c)** Rate of scratch wound closure by cardiac and lung fibroblasts in response to cytokine cocktails that broadly represent proinflammatory, anti-inflammatory and fibrotic microenvironments. Data is the mean of three biological replicates from independent experiments with SEM. **(d)** Snapshots of simulated cardiac and lung fibroblast migration with optimised modelling parameters. **(e)** Rate of gap closure by simulated cardiac and lung fibroblasts determined by the computational model. Data is the mean of 10 simulations with SD.

As immune cell-derived soluble mediators shape fibroblast activation, differentiation, and ECM remodelling in fibrosis^48–50^, cytokine cocktails were used to model proinflammatory, anti-inflammatory, and fibrotic microenvironments and assess how fibroblast phenotype influences their functional response to tissue injury. Scratch cultures were either untreated (no cytokines) or treated with defined cytokine cocktails representing distinct inflammatory states: pro-inflammatory (TNF-α, IL-1b, IL-6), anti-inflammatory (IL-10, TGFβ) or fibrotic (TGFβ, IL-4, PDGF-D). Time-lapse imaging at hourly intervals revealed fibroblasts actively migrating into the wound area over time. Untreated cardiac and lung fibroblasts migrated at similar rates (0.041 ± 0.005 mm/h and 0.044 ± 0.006 mm/h, respectively; Figure S1a), resulting in near-complete gap closure (≥95% wound area coverage) after 26.0 ± 4.4 hours and 26.0 ± 3.5 hours, respectively (Figure 2b, S1b). Cytokine stimulation did not significantly affect overall wound closure dynamics (Figure 2c, S1a,b), although there was a small, nonsignificant trend towards reduced motility in both cell types under anti-inflammatory and fibrotic conditions.

To study the impact of cell proliferation in the scratch assay context, we used cardiac and lung fibroblasts isolated from Fucci2 mice to visualise cell cycle dynamics. Fucci2 cells express red fluorescent nuclei in the G0/G1 phase and green fluorescent nuclei in the proliferative S/G2/M phase^51^. Prior to scratching, cells were arrested in G0/G1 at high confluence and with serum starvation, hence very few cells were detected in S/G2/M at the time of scratch initiation (Figure S1c,d). At 18 hours post-scratch (mid-wound closure), fibroblasts with green nuclei were observed across all conditions, indicating entry to the proliferative S/G2/M phase – likely due to the generation of free space by cell migration into the wound area. Lung fibroblasts showed a 1.5-to 2-fold higher proportion of proliferating cells (up to 26.1 ± 6.7%) compared to cardiac fibroblasts (up to 15.1 ± 3.2%) (Figure S1c,d).

An important observation was that fibroblasts did not appear to deposit collagen I during migration in the scratch assay. At 48 hours post-scratch, only faint collagen I staining was detectable around migrating fibroblasts. A distinct boundary remained where the pre-existing collagen had been removed in the process of scratching. By 96 hours, after cells repopulated the wound area, collagen I staining became uniformly distributed, and the pre- and post-scratch collagen could no longer be distinguished (Figure S2). These findings indicate that substantial collagen deposition occurs only once migration has ceased and cells have reached sufficiently high local densities. Consistent with this, in a separate assay where fibroblasts were seeded at different cell densities, collagen accumulation was observed exclusively in regions of high cell density (Figure S3a). In low-density assays fibroblasts migrated to form compact clusters, and collagen deposition became evident only within these high-density regions (Figure S3a,b). Together, these results suggest that fibroblasts may transition between migratory and matrix-depositing behaviours depending on their microenvironment.

Given the similar wound closure rates and nonsignificant impact of cytokine stimulation, we conclude that cardiac and lung fibroblasts exhibit comparable motility in the scratch assay context. Our findings indicate that fibroblast-mediated wound closure is largely driven by cell migration, with only a small contribution from proliferation such that only one-quarter of fibroblasts entered the proliferative S/G2/M phase during wound closure. Thus, whilst proliferation may support wound closure under permissive conditions, it does not appear to be a major driver. The delayed appearance of collagen fibres also supports a temporal transition between wound healing activities, whereby fibroblasts initially prioritise migration to facilitate wound closure before synthesising and depositing new collagen fibres.

### Motility drives fibroblasts to close a wound without excessive accumulation of collagen

We observed that wound closure in the scratch assay is driven primarily by fibroblast migration rather than proliferation. However, the experimental assays alone could not distinguish whether undirected (random) motility or cell-cell interactions were the dominant migratory drivers, motivating the use of a computational model to disentangle these mechanisms. In the model, undirected motility is represented by the fibroblast diffusion coefficient *D*, where we extend the previous model of Yin et al.^47^ to include decreases in motility in regions of high collagen density, as supported by observations made by Ozcelikkale et al.^52^.

Cell-cell interactions are characterised by a parameter *ϵ*, which quantifies the strength of finite-range pairwise forces encompassing both population pressure and cell-cell adhesion (Figure S4a). To assess the relative importance of undirected cell motility and cell-cell interactions to wound closure in lung and cardiac fibroblasts under different cytokine conditions, we performed systematic simulations across a range of (*D, ϵ*) values and compared the outcomes to experimental data. Consistent with experimental observations indicating that cell proliferation did not dominate wound closure, the mean cell cycle length was set to 178 hours in the computational model to ensure that wound closure simulations were primarily driven by cell migration with negligible proliferation.

We used three experimental metrics to identify the optimal (*D, ϵ*) pair that best reproduced the data. These were: (i) the time at which the wound reached 95% closure, reflecting the overall closure rate; (ii) the spatial pairwise correlation, which captures fibroblast spatial organisation, at 12, 24, and 36 hours; and (iii) collagen density within the wound area at 48 hours. The experimental values of these metrics, used as benchmarks for the computational simulations, are shown in Figure S1b,e,f. Experimentally, the control (no treatment) condition resembled the pro-inflammatory condition, while the anti-inflammatory condition resembled the fibrotic one, for both cardiac and lung fibroblasts (Figure 2c). We therefore paired these corresponding conditions and calculated the average time at which the wound reached 95% closure, using this value as the experimental benchmark for subsequent analyses. The other two experimental benchmarks (spatial pairwise cell correlation and collagen density) did not show qualitative differences across cytokine conditions and were thus averaged over all cytokine conditions. The relative differences between model predictions and experiments, in terms of these three metrics, are shown in Figures 3a-c, with contours indicating the 25% relative error threshold.

**Figure 3.**
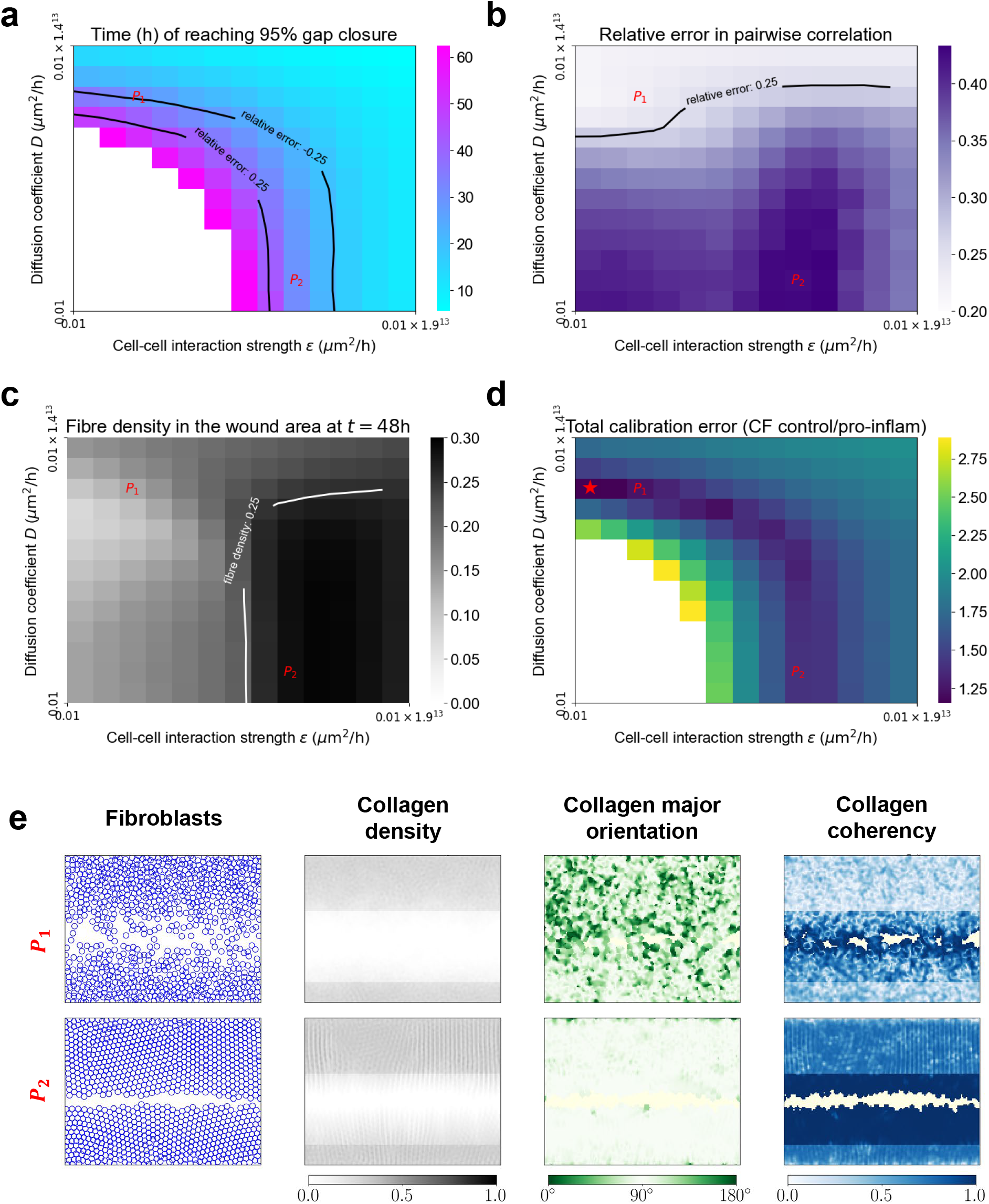
Modelling investigation of fibroblast migratory mechanisms in the scratch assay. Heatmaps visualise **(a)** the mean time to reach ≥95% gap closure, **(b)** the mean relative *L*^2^ error in the pairwise correlation of fibroblasts summed over 12, 24, and 36 hours, **(c)** the mean fibre density in the wound region at 48 hours, and **(d)** the combined total relative error from panels (a)–(c). Black contours in (a)–(c) enclose parameter values of *D* and *ϵ* yielding a relative error within 25%. The points labelled ‘*P*_1_’ (large diffusion coefficient *D*, small cell–cell interaction strength *ϵ*) and ‘*P*_2_’ (small *D*, large *ϵ*) in (a)–(c) correspond to relative errors of −0.104 and 0.104, respectively; their respective fibroblast and fibre distributions are shown in (e). The red star in (d) indicates the minimal relative error, achieved at *D* = 376.9 µm^2^/h and *ϵ* = 0.6 µm^2^/h. **(e)** Simulated cardiac fibroblast spatial patterns driven either dominantly by random motility (*P*_1_) or by pairwise cell–cell interaction (*P*_2_). Corresponding collagen fibre distributions are characterised by fibre density, major orientation, and coherency.

For cardiac fibroblasts under control and pro-inflammatory conditions, the “optimal” parameter combination – minimising the relative difference between model simulations and experimental benchmarks – is indicated by the star in Figure 3d. Optimal parameter sets for cardiac and lung fibroblasts across all conditions are shown in Figures S5a,c. Under these optimal parameters, Figures S5b,d illustrate fibroblast dynamics during wound closure and the corresponding rates of closure, where both closely match experimental observations. Overall, these results indicate that fibroblast-mediated wound closure, across both cell types and cytokine conditions, is driven primarily by undirected motility rather than cell-cell interactions. As simulation results will later illustrate, this migration mode supports efficient wound coverage whilst limiting collagen accumulation during migration and is consistent with experimental observations.

A more subtle observation is that although both undirected motility and cell-cell interactions can lead to similar wound closure rates, the spatial organisation of the assay can look very different. For example, the (*D, ϵ*) values indicated by the *P*_1_ and *P*_2_ markers in Figure 3a give similar wound closure rates, however the spatial organisation of fibroblasts differs markedly between the two. Undirected-motility-driven migration (*P*_1_) results in fibroblasts that are more dispersed within the wound area, consistent with experimental observations, whilst cell-cell-interaction-driven migration (*P*_2_) produces densely packed, cohesive fronts (Figure 3e). In the latter case, population pressure pushes fibroblasts into the wound area whilst strong cell-cell adhesion maintains tight cell packing throughout.

Moreover, collagen dynamics differ substantially between undirected-motility-driven and cell-cell-interaction-driven wound closure (Figure 3c,e), even when the baseline collagen secretion rate is low. In our computational model, fibroblasts secrete collagen aligned with their average direction of migration, with a key extension compared to Yin et al.^47^: the collagen secretion rate increases with the local cell density. This extension to the model is motivated by the experimental evidence that isolated fibroblasts secrete less collagen (Figure S3). Therefore, the undirected-motility-driven scenario (*P*_1_) leads to a more disperse fibroblast distribution, resulting in reduced ECM secretion and more isotropic alignment of the secreted collagen. This scenario is more consistent with our experimental observations. In contrast, in the cell-cell-interaction-driven scenario (*P*_2_), fibroblasts secrete denser, more aligned collagen oriented into the wound area, leading to more severe fibrotic responses.

In summary, computational modelling revealed that wound closure is driven primarily by undirected motility, rather than cell-cell interactions, across both cardiac and lung fibroblasts and all cytokine conditions. This undirected-motility-driven migration produces a dispersed fibroblast distribution while keeping collagen deposition low and weakly aligned in the wound region, consistent with our experimental findings.

### Lung fibroblasts exhibit greater proliferation and cytokine responsiveness than cardiac fibroblasts

In the experimental scratch assay, cardiac and lung fibroblasts exhibited similar migration rates; however, use of the Fucci2-expressing fibroblasts revealed that lung fibroblasts were more likely than cardiac fibroblasts to proliferate during wound closure. This observation implies that lung fibroblasts may proliferate more than cardiac fibroblasts during wound healing, and motivated us to further investigate tissue-specific differences in fibroblast proliferation using a proliferation assay.

Lung fibroblasts proliferated significantly faster than cardiac fibroblasts under all (normal and cytokine-exposed) culture conditions (Figure 4a, S6a; p<0.05). After 48 hours, only lung fibroblasts displayed cytokine-dependent changes in proliferation; proinflammatory cytokines enhanced lung fibroblast proliferation, whilst anti-inflammatory cytokines were suppressive (Figure 4a). These effects were accompanied by increased metabolic activity, as determined by the WST-1 assay, in both cardiac and lung fibroblasts exposed to proinflammatory cytokines (Figure S6b; p<0.05), consistent with the elevated energy demands of active proliferation^53^.

**Figure 4.**
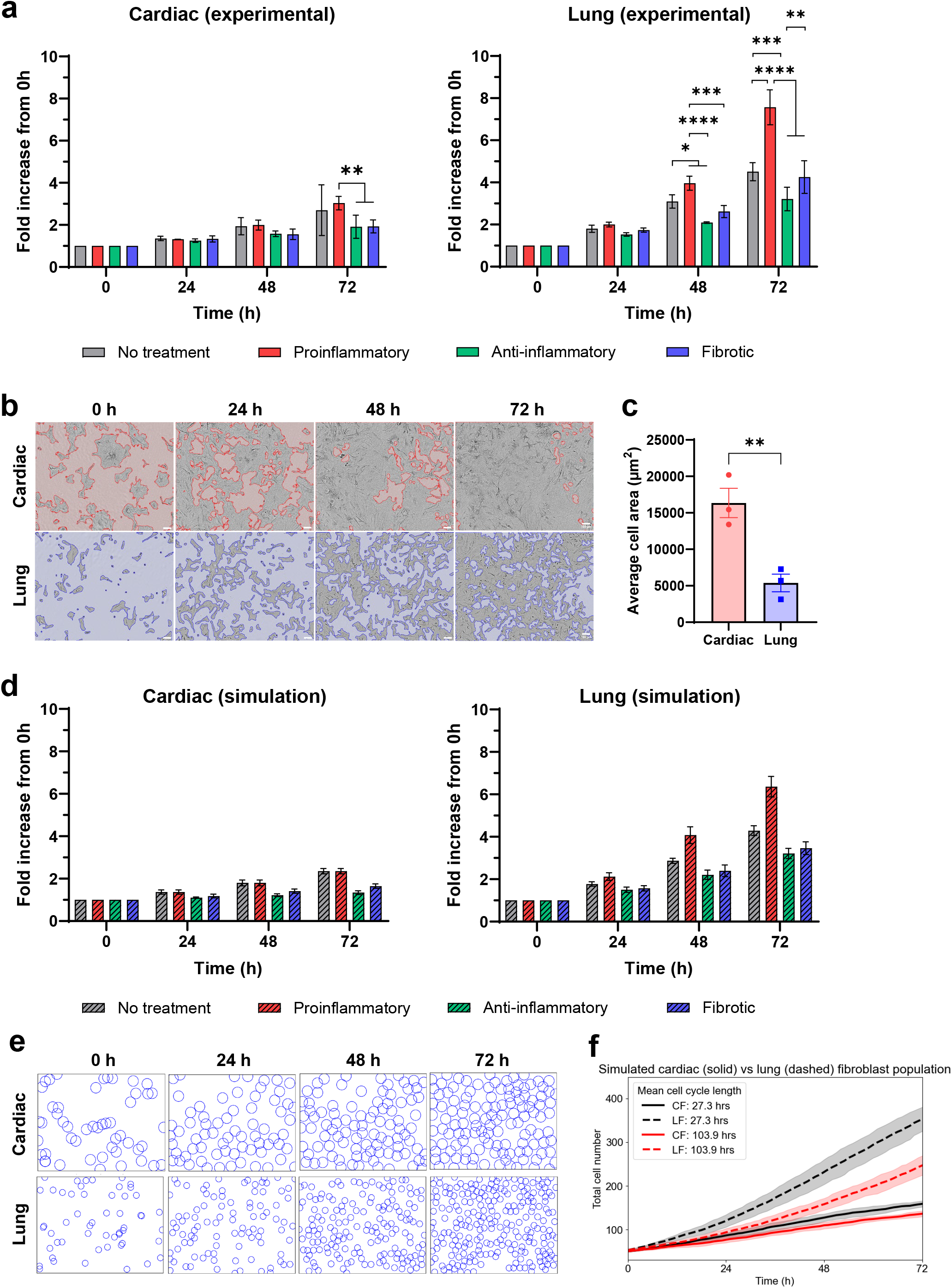
Fibroblast proliferation in response to cytokine environments. **(a)** Fold change of cardiac and lung fibroblast proliferation over time in response to different cytokine cocktails. Data is the mean of three biological replicates with SEM. Statistical significance between fibroblasts in different cytokine cocktails was determined by ANOVA with Tukey’s post-hoc testing (* *p*<0.05, ** *p*<0.01, *** *p*<0.005, **** *p*<0.001). **(b)** Snapshots of cardiac and lung fibroblasts proliferating at 24-hourly intervals. Background areas not occupied by cells are coloured in red (cardiac) and blue (lung) to distinguish the areas occupied by cells more clearly. **(c)** Average cell area size at t=0 hours. Data is the mean of three biological replicates from independent experiments with SEM. Statistical significance determined by an unpaired t-test (** *p*<0.01). **(d)** Simulated fold change of cardiac and lung fibroblast proliferation over time that best matches the experimental data. Data is the mean of ten replicates with SD. **(e)** Snapshots of simulated cardiac and lung fibroblast proliferation dynamics (control conditions) using a default cell cycle length of 47.9 hours and 42.2 hours, respectively. **(f)** Simulated cardiac and lung cell populations over time when the mean cell cycle lengths are set to either 27.3 hours (short cycle duration, plotted in black) or 103.9 hours (long cycle duration, plotted in red). The slower rate of cardiac proliferation is attributed to the larger cell areas of cardiac fibroblasts increasing the likelihood of contact inhibition. Data is the mean of ten replicates with shaded bands representing the standard deviation.

We also observed that cardiac fibroblasts modulate their cell area more dramatically than lung fibroblasts (Figure 4b,c; p<0.01). The average area of lung fibroblasts increased from 4430 ± 670 µm^2^ (high cell density in scratch assay) to 5380 ± 1210 µm^2^ (low cell density in proliferation assay, Figure 2a), whilst the average area of cardiac fibroblasts increased nearly eight-fold from 2110 ± 210 µm^2^ (high cell density in scratch assay, Figure 2a) up to 16350 ± 2010 µm^2^ (low cell density in proliferation assay). These findings suggest that cardiac fibroblasts may be more capable of dynamically regulating their surface area in response to the available space.

In the computational model, fibroblast proliferation is regulated by a parameter Δ_0_ controlling the mean cell cycle length in low density conditions, with contact inhibition of proliferation incorporated by assuming that cells divide less frequently in regions of high cell density (see Methods for more details). We used data from the proliferation assay to estimate Δ_0_ for both lung and cardiac fibroblasts with the cell motility parameters (*D, ϵ*) fixed at the values previously determined.

The estimated Δ_0_ for cardiac fibroblasts was 48 hours under control and proinflammatory conditions, increasing to 104 hours under fibrotic conditions and 178 hours under anti-inflammatory conditions (Figure S7a). In the absence of cytokines, lung fibroblasts exhibited a similar Δ_0_ (42 hours) to cardiac fibroblasts. However, culturing lung fibroblasts under proinflammatory conditions significantly decreased Δ_0_ to 28 hours, whilst fibrotic and anti-inflammatory conditions extended it to 48 and 52 hours, respectively (Figure S7b). Although lung fibroblasts proliferated faster overall, both cell types exhibited a consistent trend: anti-inflammatory and fibrotic conditions reduced proliferation rates relative to control or proinflammatory conditions. Simulated fold-changes in fibroblast population size, based on the inferred cell cycle lengths, closely matched experimental observations (Figure 4d,e).

Computational simulations also revealed that cell size influences proliferation in this assay. To highlight the impact of cell size, in simulations where Δ_0_ was set to the same value for both cardiac and lung fibroblasts, fewer cardiac fibroblasts underwent division compared to lung fibroblasts (Figure 4f). This occurs because cardiac fibroblasts are larger and so they more quickly occupy the available space, which in turn reduces cell proliferation. These results highlight that cell size and spatial constraints can significantly influence proliferation dynamics, suggesting that differences in fibroblast morphology may contribute to tissue-specific variations in wound healing and fibrosis.

### Fibroblasts exhibit organ-specific differences in collagen architecture

Collagen secretion and ECM formation are key factors in tissue fibrosis. To directly visualise collagen deposition dynamics, we utilised transgenic Col1-GFPtpz mice, which express GFP-tagged collagen Iα2 and enable live imaging of collagen secretion and assembly^54–56^. In a 72-hour collagen deposition assay, cardiac and lung fibroblasts were compared for their ability to organise collagen I, as minimal collagen III was detected (Figure S8). After 72 hours, both fibroblast types arranged collagen I into matrices with distinct architectures (Figure 5a). Consistent with previous observations (Figure 1f), cardiac fibroblasts produced diffuse collagen networks, whereas lung fibroblasts assembled more bundled fibres. The degree of fibre alignment was quantified by coherency analysis, where a value close to unity indicates highly aligned fibres, and a value close to zero indicates a disordered matrix. Lung fibroblasts were sensitive to the cytokine environment, producing more aligned collagen matrices, reflected by a greater fibre coherency, than cardiac fibroblasts under control, proinflammatory and fibrotic conditions. However, anti-inflammatory conditions markedly reduced lung collagen alignment (Figure 5b; p<0.001). In contrast, cardiac fibroblasts maintained relatively constant fibre alignment across all cytokine conditions.

**Figure 5.**
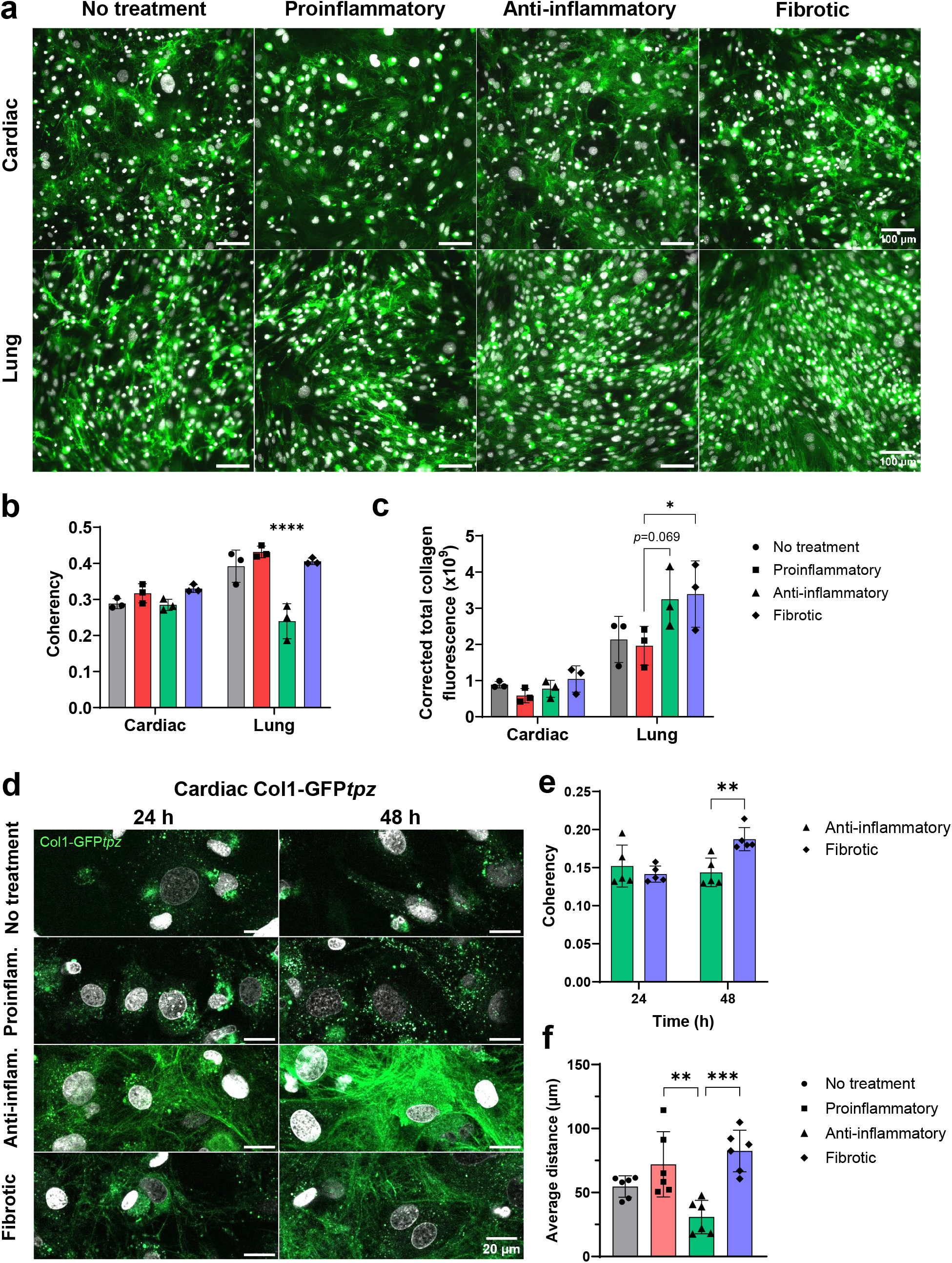
Collagen I production in response to cytokines. **(a)** Collagen I deposited by cardiac and lung fibroblasts when cultured for 72 hours in media containing ascorbic acid (100 µg/mL) and cytokine cocktails representing proinflammatory (TNF-α, IL-1β, IL-6), anti-inflammatory (IL-10, TGF-β1^lo^) and fibrotic (TGF-β1^hi^, IL-4, PDGF-D) microenvironments. **(b)** Average fibre coherency and **(c)** fluorescence intensity from collagen I matrices produced by cardiac and lung fibroblasts after 72 hours. Data is the mean of three replicates with SD. (d) Live collagen I deposition by cardiac Col1-GFP*tpz* fibroblasts at 24 and 48 hours. Nuclei stained with SPY-DNA650. **(e)** Average collagen fibre coherency at 24 and 48 hours in anti-inflammatory and fibrotic conditions. **(f)** Average distance travelled by live nuclei over 18 hours, imaging commenced after 72 hours of culture in ascorbic acid and cytokines. Data is the mean of five (e) and (f) six replicates with SD. Statistical significance determined by two-way ANOVA with Tukey’s post-hoc testing (* *p*<0.05, ** *p*<0.01, *** *p*<0.005, **** *p<*0.001).

Collagen secretion was quantified in the same collagen deposition assay by fluorescence intensity, expressed as the corrected total collagen fluorescence (CTCF). Lung fibroblasts displayed cytokine-specific variations, with anti-inflammatory and fibrotic conditions promoting greater collagen deposition compared to control or proinflammatory conditions (Figure 5c). In contrast, collagen deposition by cardiac fibroblasts did not vary significantly across the cytokine treatments (Figure 5c).

### The cell microenvironment influences extracellular matrix secretion dynamics

To understand how the early stages of matrix assembly contribute to collagen fibre organisation, primary cardiac fibroblasts isolated from Col1-GFPtpz mice were imaged at high spatial resolution. In contrast to the broader imaging scale of the 72-hour deposition assay, this approach captures collagen dynamics at the level of a few individual cells, enabling detection of subtle, cytokine-specific changes in fibre organisation during deposition.

Collagen I fibres accumulated progressively between 24 and 48 hours, but only when fibroblasts were cultured under anti-inflammatory or fibrotic conditions (Figure 5d). At 24 hours, fibre coherency was similar in both conditions, suggesting comparable initial deposition profiles (Figure 5e). However, by 48 hours high resolution imaging revealed that fibroblasts exposed to fibrotic conditions generated significantly more aligned collagen networks than those in anti-inflammatory conditions (Figure 5e; p<0.01). These findings indicate that fibroblasts begin to remodel deposited collagen into an organised architecture within the first two days of matrix formation.

Simultaneous nuclear tracking within the live collagen imaging assay revealed condition-dependent cell motility. Fibroblasts in an anti-inflammatory environment were less migratory than those in fibrotic or proinflammatory conditions (Figure 5f; p<0.01). Cells under fibrotic stimulation also appeared to deposit collagen preferentially along existing fibres, reinforcing local alignment and expanding pre-formed bundles of fibres (Supplementary Movies 1-4). These observations suggest that cell motility and local fibre guidance together shape the evolving collagen architecture.

In summary, cardiac and lung fibroblasts generated collagen matrices with distinct condition-dependent architectures: cardiac fibroblasts consistently formed diffuse collagen networks across all cytokine conditions, whereas lung fibroblasts produced more aligned matrices under proinflammatory and fibrotic conditions but showed reduced alignment under anti-inflammatory stimulation. High-resolution live-cell imaging revealed cell-level, cytokine-specific, dynamic changes in collagen organisation which may be overlooked in widefield imaging. The same imaging also showed that, although both anti-inflammatory and fibrotic conditions stimulated collagen production, fibroblast motility during deposition was reduced under anti-inflammatory conditions, which may contribute to the distinct matrix organisation observed between these conditions. These findings suggest a potential mechanistic link between cell motility and collagen alignment. To investigate this further, we employed computational modelling to examine how fibroblast motility and collagen secretion interact to shape fibre architecture and contribute to fibrotic outcomes.

### Mechanistic insights into collagen organisation via computational modelling

Motivated by previous findings that collagen matrix organisation varies with tissue type and cytokine environment, we used computational modelling as a predictive tool to investigate the interplay between fibroblast migration and collagen secretion during wound healing. We conducted numerical simulations using a “one-sided” scratch assay (Figure 6) over a long timescale (144 hours) to investigate how fibroblast motility, local cell and collagen densities, and collagen secretion rates collectively shape the ECM architecture.

**Figure 6.**
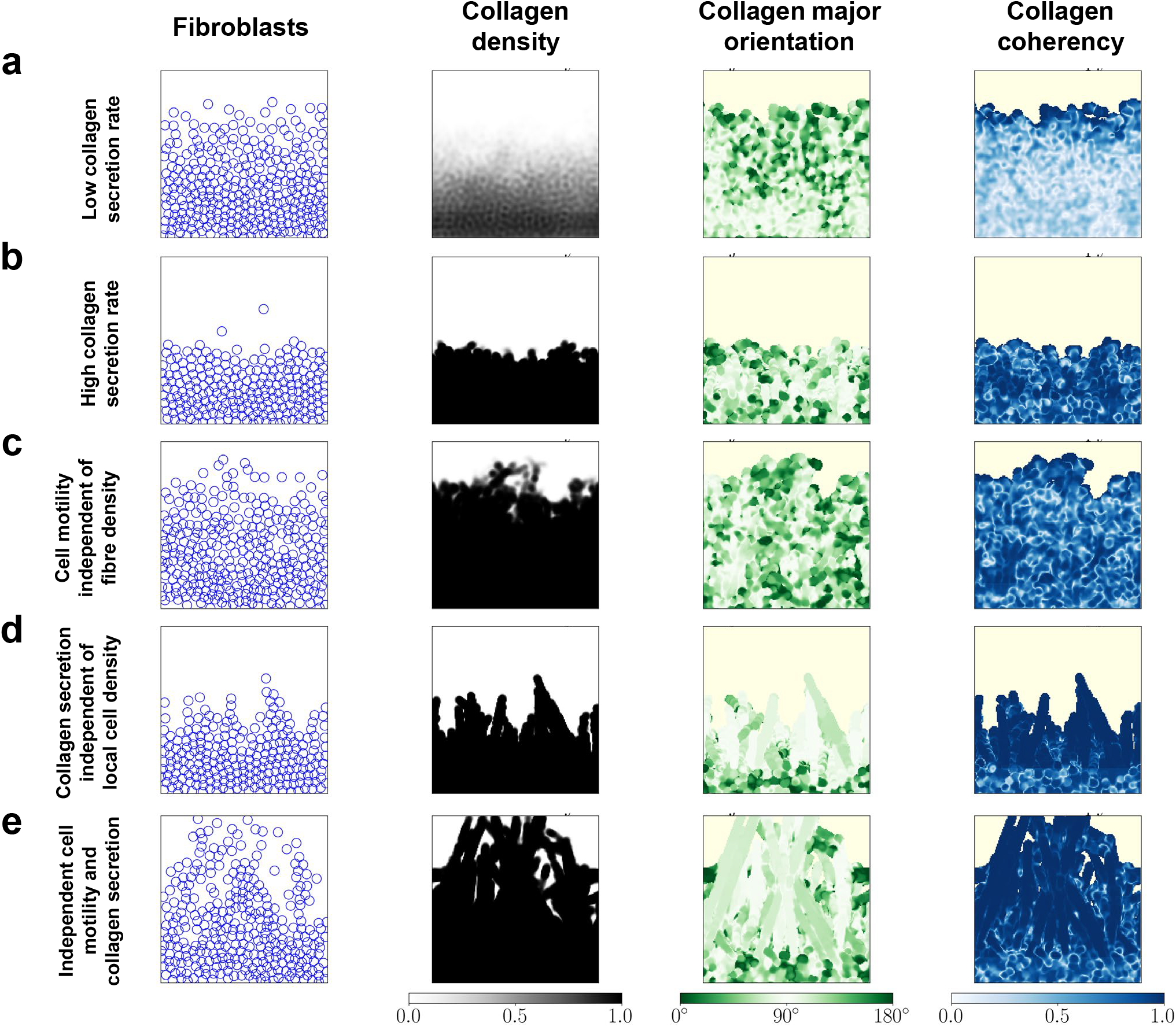
Interplay between fibroblast migration and collagen secretion during wound healing. Simulated distributions of fibroblasts and collagen fibres are shown at 144 hours under conditions of **(a)** low collagen secretion rate (s = 0.003/h) and **(b)–(e)** high collagen secretion rate (s = 0.3/h). (b) incorporates both fibre-density-dependent motility (where fibroblast random motility decreases with increasing local fibre density) and cell-density-driven collagen secretion (where collagen secretion increases with local cell density). In panels (c) and (e), the fibre-density-dependent motility assumption is removed, so fibroblast random motility remains constant regardless of fibre density. In panels (d) and (e), the cell-density-driven collagen secretion assumption is removed, resulting in a constant collagen secretion rate independent of local cell density.

We focused on two key relationships: (1) the dependence of fibroblast motility on local collagen density; and (2) the dependence of the collagen secretion rate on local cell density. In our simulations, fibroblast motility is either set constant or decreasing with increasing collagen density, and collagen secretion is either set constant or increasing with local cell density. We used the model to interrogate the interplay of these mechanisms on collagen organisation and fibroblast distributions during wound healing, under different levels of collagen secretion (high or low). The fibroblast motility parameters (*D* and *ϵ*) were fixed at the values previously inferred, and for simplicity we present results for a generic fibroblast population. Motivated by experimental observations that fibroblasts rarely proliferate during migration, we set Δ_0_ = 6000 hours so that very few cells proliferate during the simulation. Further details of the simulations can be found in Table 1.

**Table 1.** Simulated fibroblast conditions. Parameter variables for collagen secretion and fibroblast motility adjusted for each simulation in Figure 6.

| Simulation | Mechanistic assumptions |  |  |
| --- | --- | --- | --- |
|  | Collagen secretion rate | Fibre-density-dependent motility | Cell-density-driven collagen secretion |
| a | Low | Yes | Yes |
| b | High | Yes | Yes |
| c | High | No | Yes |
| d | High | Yes | No |
| e | High | No | No |

#### High collagen secretion constrains fibroblast migration and alters matrix organisation

We started by retaining the dependence of both motility on collagen density and collagen secretion on fibroblast density. Under low levels of collagen secretion, fibroblasts migrated quickly into the wound area, with relatively little collagen deposition (Figure 6a), consistent with fibroblasts cultured under proinflammatory conditions (Figures 4a, 5a-d). Increasing the levels of collagen secretion resulted in markedly reduced fibroblast migration into the wound area, and increased collagen in locally aligned but globally heterogeneous fibre networks (Figure 6b). This result was consistent with experimental observations of anti-inflammatory fibroblasts, which, in the collagen deposition assays, exhibited reduced motility and generated dense collagen networks with mixed fibre alignment (Figure 5). In contrast, removing collagen-density-dependent motility from the model resulted in markedly different migratory dynamics. Wound closure occurred more rapidly than in the collagen-density-dependent motility scenario (Figure 6c) because fibroblasts retained their ability to migrate efficiently toward the wound centre even under high levels of collagen secretion. The deposited collagen remained similarly dense, with global heterogeneity in orientation but high local alignment. These results indicate that collagen-density-dependent motility primarily regulates fibroblast migration speed (i.e. the rate of wound closure), with limited impact on collagen organisation during wound closure.

#### Cell density-dependent collagen secretion prevents fibrotic matrix formation

Removing the cell density-dependent collagen secretion assumption from the model led to the formation of dense, highly aligned collagen, with distinctive finger-like protrusions extending into the wound centre (Figure 6d). Leading-edge fibroblasts migrated into the wound centre due to population pressure from the cells behind. These front-line fibroblasts deposited collagen as they advanced into the wound and created strong contact guidance cues. Trailing fibroblasts subsequently followed and deposited collagen along these paths, reinforcing the formation of a dense, highly aligned collagen matrix. These results corresponded with our experimental observations of fibroblasts under fibrotic conditions, which generated dense, highly aligned fibres and retained high levels of motility during collagen deposition (Figures 5b,c,e,f). When fibroblast motility was independent of collagen density *and* collagen secretion was constant regardless of the local cell density, fibroblast migration and wound closure were further accelerated (Figure 6e). However, the resulting matrix organisation remained similarly dense with highly aligned collagen forming across the wound domain. The disproportionate accumulation of collagen-rich matrices and reduced cellular content mimicked the formation of a maladaptive fibrotic tissue, which reflects an abnormal wound healing response.

In conclusion, these results suggest that accelerated wound closure alone is insufficient to prevent fibrotic tissue formation. Rather, reparative wound healing requires coordinated feedback between collagen-density-dependent fibroblast migration and cell-density-dependent collagen secretion. Disrupting these feedbacks can lead to rapid wound closure accompanied by the dense, highly aligned collagen networks that are characteristic of pathological fibrosis.

## Discussion

In this study, we compared phenotypically distinct cardiac and lung fibroblasts as drivers of cardiac and pulmonary fibrosis, revealing tissue-specific differences in cell morphology, functional behaviour, and collagen I deposition. Lung fibroblasts maintained a consistent morphology and size regardless of their environment; however, cardiac fibroblasts displayed irregular morphologies and density-dependent morphological plasticity, dynamically altering their cell area and shape. This plasticity exhibited by cardiac fibroblasts may reflect regulation of the cytoskeleton via mechanical pathways, as reported by Zhou et al.^57^, who observed similar density-dependent changes in mouse embryonic fibroblasts.

Mechanical pathways are also recognised as key drivers of fibroblast-to-myofibroblast differentiation^58^. A phenotypic indicator of this transition is expression of α-SMA which represents the incorporation of actin into stress fibres, facilitating tissue repair and fibrosis by regulating cell stiffness and extracellular matrix contraction^8,9^. Lung fibroblasts expressed higher levels of α-SMA than cardiac fibroblasts, indicating a greater tendency toward myofibroblast activation. Previous reports attributed the difference in α-SMA to a lower β3 integrin expression in lung fibroblasts compared to cardiac fibroblasts, which regulates wound healing, fibroblast phenotype and TGF-β signalling^59^. In our study, lung fibroblasts also formed collagen matrices with more distinct collagen fibre bundling than matrices produced by cardiac fibroblasts. In dermal fibrosis, α-SMA-positive fibroblasts were detected along aligned collagen bundles^60^, and aligned collagen fibrils can upregulate α-SMA expression *in vitro*^61^. Although derived from dermal models, these findings support the possibility that increased α-SMA in lung fibroblasts may both drive and reinforce collagen bundling through a positive feedback loop. Together, these results suggest that tissue origin contributes to differences in fibroblast morphology and activation.

To assess behavioural differences between tissue-specific fibroblasts, a series of functional assays were performed. Scratch assays showed that cardiac and lung fibroblasts migrate at comparable rates and deposit minimal collagen during gap closure. This suggests that core migratory mechanisms are conserved across organs in the early stages of tissue injury. Substantial collagen deposition became evident only after migration ceased and cells reached sufficiently high local densities, indicating behavioural plasticity and a phenotypic switch between migratory and ECM synthesis states^62^. Consistent with this, fibroblasts seeded at low density migrated to form densely populated regions, which appeared to be a prerequisite for collagen deposition. Analogous to the “go-or-grow” hypothesis describing reversible transitions between migration and proliferation phenotypes^63^, fibroblasts seem to switch between migration and ECM-producing phenotypes in response to their microenvironmental cues. This switching aligns with the temporal sequence of tissue repair, in which matrix deposition follows cellular migration^64^, a dynamic that is also reproduced in computational models of tissue remodelling^65^.

The complex interplay between cellular and tissue-level factors makes it difficult to identify targetable pathways for modulating fibrosis. To address this, we developed a bespoke computational model based on Yin et al.^47^ to investigate the mechanisms underlying the fibroblast behaviours observed in our *in vitro* assays and gain insight into the cellular factors driving fibrotic outcomes. By comparing model predictions with our experimental scratch assay data, we identified that the primary driver of wound closure was undirected cell motility rather than cell-cell interactions (driven by population pressure and adhesion). Computational simulations revealed that undirected motility leads to a more dispersed fibroblast distribution within the wound area and minimal collagen deposition, consistent with experiments. The model also predicted that collagen formed under this regime is globally heterogeneous in orientation and less locally aligned than when migration is driven by cell-cell interactions.

Proliferation assays confirmed that lung fibroblast populations proliferate faster than cardiac fibroblasts, reflecting a greater demand for lung tissue to require constant repair and cell turn-over due to continuous exposure to environmental pathogens, debris and airborne insults^66,67^. Both fibroblast types exhibited increased proliferation and metabolic activity under proinflammatory stimulation, reflecting a coordinated and context-dependent response to tissue injury^68^.

Collagen deposition and organisation are key features of organ fibrosis. Cardiac fibrosis is characterised by the accumulation of highly aligned collagen following acute injury such as myocardial infarction, which is conserved across species^69,70^. Pulmonary fibrosis also presents with increased collagen density and alignment^19,71–73^, but reports of collagen organisation vary. In bleomycin-induced mouse models, fibrosis produced linear, aligned collagen over time^71^, whereas human lung tissue with idiopathic pulmonary fibrosis showed densely accumulated wavy fibres compared with the sparser, more linear architecture in healthy lungs^19,73^. Our collagen I deposition assays revealed distinct matrix architectures between cardiac and lung fibroblasts, likely reflecting tissue-specific demands. Cardiac fibroblasts formed diffuse, basket-weave collagen networks with frequent interconnecting fibres, which did not vary significantly under different cytokine cocktails. In contrast, lung fibroblasts produced defined collagen bundles, which varied in alignment under different cytokine environments. Under control, proinflammatory and fibrotic conditions, lung collagen matrices were more aligned than the corresponding cardiac matrices, whereas anti-inflammatory conditions reduced lung collagen alignment to that observed in cardiac matrices. Cardiac mechanical strain is a major determinant of collagen alignment in the heart at baseline and following myocardial infarction^65,70,74^. The absence of strain *in vitro* may have led to the heterogeneous collagen organisation by cardiac fibroblasts observed across all cytokine conditions, suggesting that biochemical cues are insufficient to compensate for mechanical loading and strain within the heart. Whilst lung fibroblasts also experience cyclic stretch from respiration, the amplitude and frequency of these forces differ from myocardial contractility^75–77^. Cardiac fibroblasts have been reported to tolerate stiffer substrates without adopting a myofibroblast phenotype (e.g. 50 kPa *in vitro*), whereas lung fibroblasts activated into myofibroblasts on softer substrates (∼20 kPa)^78^, suggesting a greater tolerance of stiffer environments by cardiac fibroblasts. Consistent with this, TGF-β, which has a key role in fibrosis, increased COL1A expression by ∼15-fold in lung fibroblasts but only ∼1.15-fold in cardiac fibroblasts^59^, providing further evidence of tissue-specific biochemical sensitivities. Together, these intrinsic mechanical and biochemical differences likely contribute to the distinct progression of cardiac and pulmonary fibrosis observed clinically^27^.

As widefield imaging revealed no differences in cardiac collagen organisation across different cytokine conditions, we employed high-resolution live imaging of GFP-tagged collagen produced by transgenic cardiac fibroblasts (Col1-GFPtpz) to capture early collagen deposition. Under anti-inflammatory conditions, cardiac matrices exhibited mixed fibre orientations, resembling the basket-weave architecture of resolving scars^79,80^, whereas fibrotic conditions promoted more collagen alignment, characteristic of pathological scarring^81,82^. Fibroblast motility also differed, with fibrotic fibroblasts migrating further, consistent with findings that fibroblasts migrate faster on highly aligned topographies^83^. As collagen architecture influences cell motility^52^, these results suggest that fibroblast phenotype regulates both cell migration and matrix organisation, establishing a feedback loop between motility and collagen alignment.

We used the computational model as a predictive tool to extend beyond our experimental findings and examine how fibroblast motility shapes collagen organisation. We focused on two relationships: (1) the dependence of fibroblast random motility on local collagen density; and (2) the dependence of collagen secretion rate on local cell density. Simulations showed that when fibroblast motility decreases with increasing collagen density, a low-collagen-secretion regime resembled pro-inflammatory fibroblasts behaviours, with limited collagen deposition and slightly faster wound closure. In contrast, a high-collagen-secretion regime resembled anti-inflammatory fibroblast behaviours where increased collagen deposition restricted migration. These simulation results provide a mechanistic interpretation of the experimentally observed switch between migratory and ECM-producing fibroblast phenotypes. By allowing motility to decrease with increasing collagen density and collagen secretion to rise with local cell density, we have shown that fibroblasts prioritise migration in low-collagen regions to facilitate wound closure and then transition to matrix production as local crowding and collagen accumulation increase. Simulations further showed that when collagen secretion increases with the local cell density, the formation of highly aligned, finger-like collagen protrusions is suppressed, thereby reducing the likelihood fibrotic matrix organisation. In contrast, when collagen secretion and fibroblast motility are independent of the local microenvironment simulations indicate that chronic healing defects are prevented but at the cost of fibrotic matrix accumulation.

Overall, our study demonstrates that both tissue origin and phenotype shape key fibroblast behaviours – migration, proliferation, and matrix deposition – ultimately influencing collagen organisation and fibrosis. Integrating experimental and computational approaches revealed mechanistic links between fibroblast motility and ECM architecture, highlighting motility as a potential therapeutic target. A deeper understanding of shared and tissue-specific fibroblast mechanisms may guide the development of both broad-spectrum antifibrotic strategies targeting core pathways and organ-specific therapeutics tailored to individual tissue biology. Future studies incorporating additional cell types or mechanical cues may further elucidate fibroblast phenotypic transitions and support the adaptation of the computational model to simulate therapeutic interventions.

## Materials and methods

### Experimental methods

#### Animals

Wild-type C57BL/6 mice were obtained from Charles River Laboratories (UK). Fucci2 mice (*Tg(Gt(ROSA)26Sor-Fucci2)#Sia*) were kindly provided by Professor Shankar Srinivas (University of Oxford). GFP*topaz*-collagen1α2 (Col1-GFP*tpz*) mice were generated and donated by Professor Sarah Dallas (University of Missouri-Kansas City). Transgenic mouse lines were maintained as heterozygotes by breeding with C57BL/6 mice.

Mice were housed and maintained in a controlled environment by the University of Oxford Biomedical Services. Both male and female adult mice (8-to 16-week-old) were used in experiments. Breeding and culling by Schedule 1 procedures were performed in accordance with the UK Animals (Scientific Procedures) Act 1986. Procedures were covered under UK Home Office Project Licence PP3194787 and approved by the University of Oxford Animal Welfare and Ethical Review Body (AWERB).

#### Isolation of primary fibroblasts

Murine hearts and lungs were harvested and placed into ice-cold PBS. Under sterile conditions, tissues were minced into small pieces (1-2 mm) and suspended in HBSS containing Liberase Thermolysin Medium (Roche; 0.1 units/mL) and DNase I (50 units/mL). Tissue was transferred into gentleMACs C Tubes (Miltenyi Biotec) and digested using the gentleMACS Octo Dissociator (Miltenyi Biotec). Cardiac and lung tissues were digested using program settings 37C_Multi_G and 37C_m_LDK_1, respectively. DMEM media containing 20% (v/v) FBS was added to digested suspensions and filtered through a 70 µm cell strainer. Cell suspensions were centrifuged (300 x *g*) for 5 minutes and supernatant aspirated. Debris removal was performed using the Debris Removal Solution (Miltenyi Biotec) according to the manufacturer’s instructions.

Isolated cells were resuspended in complete growth medium consisting of DMEM/F-12 (Sigma-Aldrich) supplemented with 10% (v/v) FBS (Gibco), L-glutamine (2 mM) and 100 units/100 µg/mL penicillin/streptomycin (Sigma-Aldrich). Cells isolated from a single organ were plated into one well of a 6-well plate (10 cm^2^/single organ) and scaled accordingly if hearts or lungs were pooled together. Media was removed and cells gently washed with PBS prior to adding fresh media the following day. Media was replaced every 2-3 days until 90% confluence was attained 5-7 days following isolation. Cells were used for experimental purposes up to the third passage, after which cultures were discontinued. Successful isolation of cardiac and lung fibroblasts was confirmed by positive expression of vimentin, CD90 and platelet-derived growth factor alpha (PDGFR□), and negative expression of the endothelial marker CD31 (Figure S9).

#### Cytokine stimulation

Fibroblasts were cultured in cytokine cocktails to mimic biochemical environments associated with inflammation and wound healing. Proinflammatory: tumour necrosis factor (TNF)-□ (20 ng/mL), interleukin (IL)-1β (10 ng/mL) and IL-6 (50 ng/mL); anti-inflammatory: IL-10 (10 ng/mL) and transforming growth factor (TGF)-β (2 ng/mL); fibrotic: TGF-β (10 ng/mL), IL-4 (10 ng/mL) and platelet-derived growth factor (PDGF)-D (200 ng/mL). TNF-□, IL-1β, IL-4, IL-6 and IL-10 were sourced from PeproTech, whilst TGF-β and PDGF-D were purchased from Bio-Techne and R&D Systems, respectively.

#### Immunofluorescence

Fibroblasts were cultured on gelatin-coated (300 µg/cm^2^) Ibidi 8-well slides and fixed in 4% paraformaldehyde (PFA) in PBS for 15 minutes at room temperature. Cells were washed three times with PBS and permeabilised by 0.1% (w/v) Triton X-100 in PBS for 15 minutes at room temperature. Non-specific binding was blocked using 3% (v/v) normal goat serum and 5% (w/v) bovine serum albumin (BSA) in PBS (blocking buffer) for 1 hour at room temperature. Primary antibodies were diluted (1:200) in blocking buffer and incubated for 1 hour at room temperature or overnight at 4 °C. Cells were washed three times with PBS and followed by addition of species-specific secondary antibodies diluted (1:500) in blocking buffer for 1 hour at room temperature and protected from light. If required, DAPI and Phalloidin-FITC were added to the secondary antibody mixture. Cells were washed three times with PBS and kept in PBS for short-term storage and imaging. Imaging was performed using a Leica DMi8 inverted microscope or a Zeiss 980 LSM confocal microscope.

Primary antibodies included anti-□-SMA (PA5-85070, Invitrogen), anti-CD31 (558736, BD Pharmingen), anti-CD90 (ab307736, Abcam), anti-collagen I (ab21286, Abcam), anti-collagen III (ab218164, Abcam) anti-GFP (ab13970, Abcam), anti-PDGFR□ (PA5-34739, Invitrogen), anti-vimentin (ab92547, Abcam). Secondary antibodies sourced from Invitrogen included donkey anti-rabbit IgG Alexa Fluor 546, goat anti-rabbit IgG Alexa Fluor 594, goat anti-rat IgG Alexa Fluor 647, and goat anti-chicken IgY Alexa Fluor 647.

#### Scratch assay

Wells of an Incucyte® Imagelock 96-well Microplate (Sartorius) were coated in gelatin (300 µg/cm^2^; Stemcell Technologies) for 30 minutes at room temperature. Cells were seeded at a density of 5,000 cardiac fibroblasts/well or 8,000 lung fibroblasts/well. Lung cells appeared smaller and required a higher initial cell density to achieve confluence at the same time as cardiac fibroblasts. Upon confluence, complete growth medium was replaced with low-serum media consisting of DMEM/F-12 supplemented with 0.5% (v/v) FBS and 100 units/100 µg/mL penicillin/streptomycin for 24 hours. Scratches in confluent cell monolayers were made using the Incucyte® Woundmaker 96-Tool (Sartorius) and cell debris was removed by washing cells with PBS twice. Complete growth medium containing 10% (v/v) FBS and cytokine cocktails were added to scratched cells and a SPY595-DNA probe (1:1000 dilution, Spirochrome) was added to visualise nuclei. Cells were incubated in the Incucyte® S3 Live-Cell Analysis System (Sartorius) and imaged at hourly intervals for 48 hours. Five technical replicates were set-up for each condition and repeated three times in independent experiments using different biological specimens.

#### Proliferation assay

Wells of a 96-well plate were coated in gelatin (300 µg/cm^2^; Stemcell Technologies) for 30 minutes at room temperature. Cells were seeded at a density of 2,500 cardiac fibroblasts/well or 3,000 lung fibroblasts/well and left to adhere for 24 hours. Fresh medium containing cytokine cocktails was added to the cells. Cells were incubated in the Incucyte® S3 Live-Cell Analysis System (Sartorius) and imaged at 24-hour intervals for three days. Four technical replicates were set-up for each condition and four images acquired per well with the most representative image used for analysis. Experiments were repeated using three biologically independent specimens. Cell counting of brightfield images was performed manually using FIJI (v1.54p)^84^.

#### Collagen matrix production and imaging

Cells were seeded into gelatin-coated (300 µg/cm^2^) Ibidi 8-well slides at 12,000 cardiac fibroblasts/well and 15,000 lung fibroblasts/well. Cells were left to adhere for 24 hours and the media replaced with growth medium containing ascorbic acid (50 µg/mL) with or without cytokines. Media was replaced daily for up to 7 days. Cells were either imaged live or fixed in 4% (w/v) PFA for antibody staining. Cell nuclei for live cell imaging were dyed with a SPY650-DNA probe (1:1000 dilution; Spirochrome) two hours prior to imaging. Imaging was performed using Zeiss 880 and 980 LSM confocal microscopes equipped with cell incubation systems.

Collagen matrices generated over 7 days (Figure 1) were decellularised by lysing the cells in 0.1% (w/v) sodium dodecyl sulphate and DNase I (50 units/mL) for 5 minutes. Matrices were gently washed with PBS three times prior to fixing in 4% (w/v) PFA.

#### Image analysis

All image analyses from experimental results were conducted in FIJI (v1.54p)^84^. Cell and collagen orientation and coherency heat maps were produced using the “Analysis” and “Distribution” functions in the FIJI plugin OrientationJ^85,86^. The “Measure” function in OrientationJ^87^ was employed to determine the major orientation angle using the direction of F-actin filaments in the cytoskeleton of each cell. The nematic order parameter (S) represents the influence of the local cell population on individual cell alignment and was calculated by 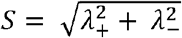, where *λ*_±_ are the eigenvalues of the nematic tensor *Q*:

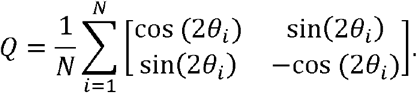

of a total of cells.

Here, *θ*_*i*_ is the major orientation angle with respect to the horizontal *x*-axis of the *i*th cell, out of a total of cells.

Expression of □-SMA stress fibres in myofibroblasts was differentiated from basal □-SMA expression in fibroblasts by manually counting □-SMA-positive cells that were observed to possess distinct stress fibres and using the FIJI plugin JACoP for colocalisation image analysis^88^. The Manders’ colocalisation index was calculated by analysing the degree of overlap of F-actin fluorescence with □-SMA fluorescence. A value of unity indicates that □-SMA expression is completely colocalised with F-actin filaments due to the formation of contractile □-SMA stress fibres.

Gap closure in the scratch assay and cell area measurements in the proliferation assay were quantified using in-house segmentation macros. These macros can be found in the supplementary attachments.

Collagen orientation and coherency was determined using the FIJI plugin OrientationJ “Vector Field” function^85,86^ with post-processing to remove background values. The Harris Index of collagen matrices was determined using the OrientationJ “Corner Harris” function. Corrected total collagen fluorescence (CTCF) intensity was calculated by

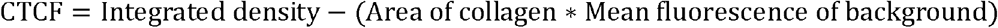

Nuclei tracking in the live collagen assay was performed using the FIJI plugins Stardist^89^ and TrackMate^90,91^.

### Mathematical framework

Our two-dimensional mathematical model builds upon the framework developed by Yin et al.^47^ and describes the interactions between two key components: cells and collagen. In this model, cells are represented as discrete agents that migrate over a continuous collagen field. The original model includes several core mechanisms: constant random motility and cell-cell interactions, both modulated by collagen alignment via contact guidance; cell proliferation; and constant collagen secretion and degradation. We retain these mechanisms but introduce two biologically motivated innovations based on our experimental findings: (1) random cell motility depends on local collagen density; and (2) collagen secretion is regulated by local cell density. In the following sections, we first describe how cells and collagen are represented and then detail the dynamics of each component. Python codes to simulate the model are available at https://github.com/YuanYIN99/OrganSpecificFibrosis.git.

#### Representations for cells and collagen

Following Yin et al.^47^, cells are represented as point particles whose positions at time *t* are given by ***X***^*i*^(*t*) ∈ ***G*** ⊆ ℝ^2^, where *i* = 1, …,*N*(*t*), and *N*(*t*) denotes the total number of cells in the domain ***G*** at time *t*. The collagen distribution is described by a continuous tensor field **Ω** (***x***,*t*) whose diagonalised form is

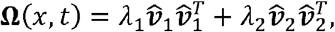

where 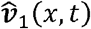 and 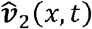 are orthonormal eigenvectors corresponding to the major and minor collagen orientations, respectively. The associated eigenvalues *λ*_1_(*x,t*) ≥ *λ*_*2*_ (*x,t*) ∈ [0,1] characterise the collagen densities along these directions. Given such definition of **Ω**, one can extract both the total collagen density, given by *λ*_1_(*x,t*) + *λ*_*2*_ (*x,t*) ∈ [0,1], and the collagen coherency, defined as (*λ*_1_ − *λ*_2_)/(*λ*_1_ + *λ*_2_) ∈ [0,1]. The coherency, as defined in the collagen fibre image analysis tool OrientationJ, measures the degree of alignment of collagen: a value of zero corresponds to an isotropic distribution, while a value of one indicates perfect alignment along the major collagen direction 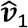. Figure S4b provides a schematic diagram of the model representation of cells and collagen.

#### Cell dynamics

Following Yin et al.^47^, the motion of each cell *i* = 1,…,*N*(*t*) is governed by the equation

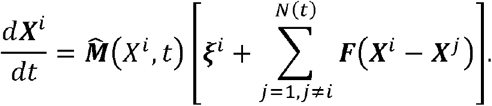

In this formulation, ***ξ***^*i*^ represents random cell motility,*F*(***X***^*i*^ − ***X***^*j*^) captures the pairwise interaction force between cell *i* and cell *j*, and 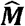 is a normalised length-preserving transformation matrix encoding contact guidance that re-orients cell according to the major collagen orientation imposed by the local collagen distribution. Each of these components is detailed below.

##### Collagen-dependent random motility

We assume each cell *i* undergoes random motion driven by a 2-by-1 stochastic force ***ξ***^*i*^, sampled from a Gaussian distribution with zero mean. However, rather than using a constant variance of 2D/Δ*t*, we extend the model of Yin et al.^47^, such that the variance becomes 2*D*_dff_ (*λ*_1_ *+ λ*_2_)/Δ*t*,, where *D*_dff_ (*λ*_1_ *+ λ*_2_) is the effective diffusion coefficient, modelled as a function of the total collagen density (*λ*_1_ *+ λ*_2_) at cell *i*’s location ***X***^*i*^(*t*), and is the fixed numerical time step. Motivated by our experimental observations that cells have lower motility in collagen-rich regions (where *λ*_1_ *+ λ*_2_ ≈ 1), we choose a simple linear relationship:

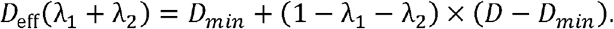

Here *D*_*min*_ ≥ 0 is the minimal diffusion coefficient in collagen-saturated regions whereas *D* > *D*_*min*_ > 0 represents the baseline diffusion coefficient in collagen-free areas.

##### Pairwise cell-cell interactions

We adopt the same formulation as in Yin et al.^47^ where the interaction force experienced by cell *i* from cell *j* arises from a combination of cell-cell adhesion and volume exclusion, and is modelled as a 2-by-1 vector

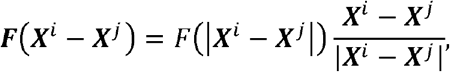

where the scalar function *F*(|***X***^*i*^ − ***X***^*j*^|) defines the magnitude of the force based on the distance |***X***^*i*^ − ***X***^*j*^|. The force magnitude is derived from a Lennard-Jones potential where

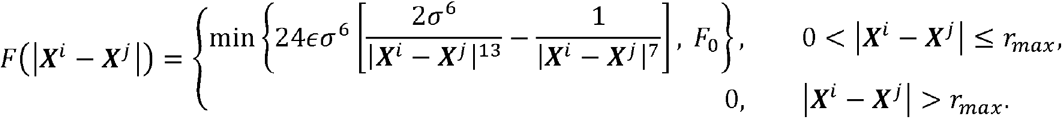

Here σ > 0 is a measure of the cell diameter, and ϵ > 0 regulates the strength of pairwise interactions. The cutoff *F*_0_ > 0 ensures the force remains bounded as *r* → 0 and, *r*_max_ > 0 defines the maximum interaction range between two cells.

##### Contact guidance

To model how the local collagen distribution influences the migratory direction of cell *i*, we follow the framework introduced by Yin et al.^47^, in which both the stochastic and the cell-cell interaction forces are re-oriented by a normalised transformation matrix 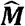, where

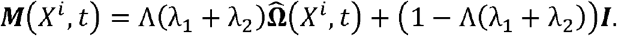

Here, 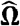 is the normalised form of the tensorial representation of collagen **Ω**, and ***I*** is the 2-by-2 identity matrix. The function Λ(*λ*_1_ *+ λ*_2_) ∈ [0,1] controls the strength of contact guidance and depends on the local collagen density. It is defined as

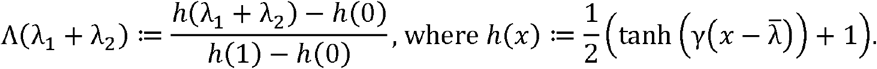

Here, 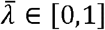 represents the critical collagen density at which the strength of contact guidance is half-maximal, and *γ* >0 controls the steepness of the transition around 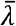. In regions where collagen is both dense (*λ*_1_ *+ λ*_2_ ≈ = 1) and highly aligned (*λ*_1_ − *λ*_2_)/ (*λ*_1_ *+ λ*_2_) ≈ = 1), 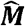 strongly re-orients towards the major collagen orientation 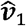. In contrast, in regions of low or isotropic collagen density, the influence of contact guidance diminishes.

##### Cell proliferation

We adopt the proliferation model from Yin et al.^47^ without modification. In this model, the probability *P*_*p*_ that a cell divides during the time interval [*t, t* + Δ*t*) decreases with increasing local cell density, reflecting space limitation due to crowding, with

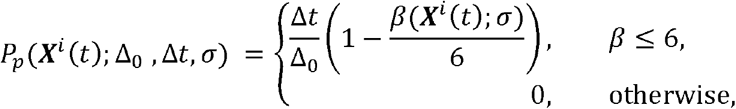

where Δ_0_ > 0 is the mean cell cycle length in the absence of crowding, and *β*(***X***^*i*^ (*t*); *σ*) denotes the number of nearest neighbouring cells ‘sensed’ by cell *i* at position ***X***^*i*^ (*t*). Nearest neighbours are defined as cells located within the repulsive interaction range, which is proportional to (a measure of cell diameter). The value six corresponds to the assumed maximum packing number in two dimensions^92^. Accordingly, *β*(***X***^*i*^ (*t*); *σ*) /6 serves as our measure of local crowdedness.

#### Collagen dynamics

We assume that collagen dynamics are governed by degradation and secretion by cells, represented by the equation

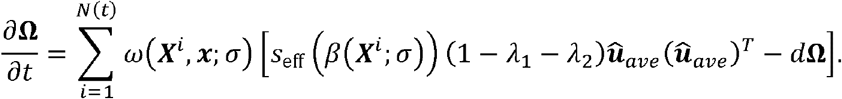

Here, the function *ω* (*X*^*i*^, *x*; *σ*) denotes the local influence of cell *i* on the collagen field, and is defined as

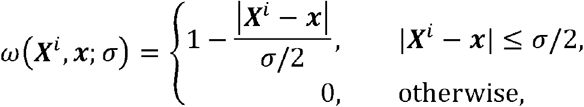

following Yin et al.^47^. This compactly supported function localises each cell’s effect within a radius of 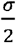 with the effect decreasing linearly from the cell’s centre. The term *d* > 0 represents the uniform degradation rate of collagen of all orientations. Collagen secretion occurs along the average direction of cell i’s recent movement 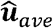, where the average velocity over [*t* − *m, t*,] ***u***_*ave*_(***X***^*i*^, *t*;*m*) is defined as

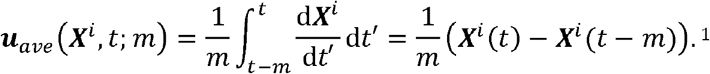

The amount of collagen secreted is modulated by the factor (1 − *λ*_1_ − *λ*_1_), which ensures that no additional collagen is deposited in regions already saturated with collagen. Importantly, unlike Yin et al.^47^, who assumed a constant secretion rate, we extend the secretion mechanism based on experimental observations indicating that fibroblasts tend to secrete more collagen when other cells are present nearby. Specifically, we define the effective secretion rate *S*_eff_ as

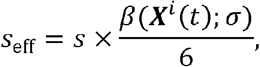

where *S* > 0 is the default secretion rate when a cell is surrounded by six neighbours. Recall that counts the *β*(***X***^*i*^, *x*; *σ*) number of nearest neighbouring cells (within the repulsive interaction range), and *β*(***X***^*i*^, *x*; *σ*) /6 is a measure of local cell density. Thus, this formulation introduces a simple linear relationship between local cell density and collagen secretion.

## Acknowledgments

YY is funded by the Engineering and Physical Sciences Research Council (EP/W524311/1). This work was supported by a grant from the Simons Foundation (MP-SIP-00001828, REB) and the University of Oxford’s John Fell Fund. We thank the Oxford-ZEISS Imaging Facility for use of the Zeiss 980 LSM confocal microscope. We thank Professor Shankar Srinivas (University of Oxford) for use of Fucci2 mice and the Zeiss 880 LSM confocal microscope. We thank Professor Sarah Dallas (University of Missouri-Kansas City) for the GFP*topaz*-collagen1α2 mouse line. We thank Professor Daniel Ebner (University of Oxford) for use of the Incucyte Woundmaker tool and Target Discovery Institute facilities. For the purpose of open access, the authors have applied a CC BY public copyright licence to any author accepted manuscript arising from this submission.

## Declaration of interests

The authors declare no competing interests.

## Supplementary Information

### Supplementary Methods

#### Metabolic assay

Wells of a 96-well plate were coated in gelatin (300 µg/cm^2^; Stemcell Technologies) for 30 minutes at room temperature. Cardiac and lung fibroblasts were seeded at a density of 4,000 cells/well and left to adhere for 24 hours. Medium was replaced for fresh containing cytokine cocktails (200 µL/well) and cultured for a further 24 hours. Cell Proliferation Reagent WST-1 (Roche) was added to the cell culture medium (20 µL/well) and incubated at 37 °C for 4 hours. The production of soluble formazan dye was measured at absorbance 450 nm with a plate reader.

### Computational set-up for assays

#### Scratch assay

To provide mechanistic insights for the experimental results about scratch assays, we initialise our model as shown in Figure S4c, where the black box encloses the field of view (FOV) of the same size as in the experiments, and the red box encloses the initial wound region of the size averaged from the experimental data. In the FOV outside the wound area, fibroblasts are initialised to form confluent monolayers, where the average cell numbers are obtained from the experimental data. Outside the FOV are what we call ‘cell pools’ for simulation purposes, to model the vertical in-flux of fibroblasts into the FOV during the wound closure process. We assume periodic x-boundary conditions for fibroblasts. Everywhere in the simulation domain except for the wound region, we initialise with sparse uniformly distributed collagen of density 0.1 in random directions to mimic the experiments.

#### Proliferation assay

To provide mechanistic insights for the experimental results about proliferation assays, we initialise our model as shown in Figure S4d, where fibroblasts are distributed uniformly into the simulation domain, and the domain size matches that of the experimental FOV. The number of fibroblasts found by averaging the experimental initial cell populations among replicates. For fibroblasts, we assume periodic boundary conditions in both x- and y-directions. We assume no collagen is present at the start of the proliferation assay simulations.

#### One-sided scratch assay

To predict fibrotic outcomes and the architecture of fibroblast-deposited collagen, we initialised our model as illustrated in Figure S4e. The black box denotes the FOV with size [0, 1600] × [0, 1600] *μ*m^2^, while the red box marks the initial wound region spanning [0, 1600] × [0, 1330] *μ*m^2^. Within the FOV but outside the wound area, 60 fibroblasts were initialised to form a confluent monolayer. Beyond the FOV, 66 fibroblasts were initialised in the ‘cell pool’ to maintain confluence. Periodic boundary conditions were applied along the x-axis. Everywhere in the simulation domain except for the wound region, we initialise with sparse uniformly distributed collagen of density 0.1 in random directions.

**Figure S1.**
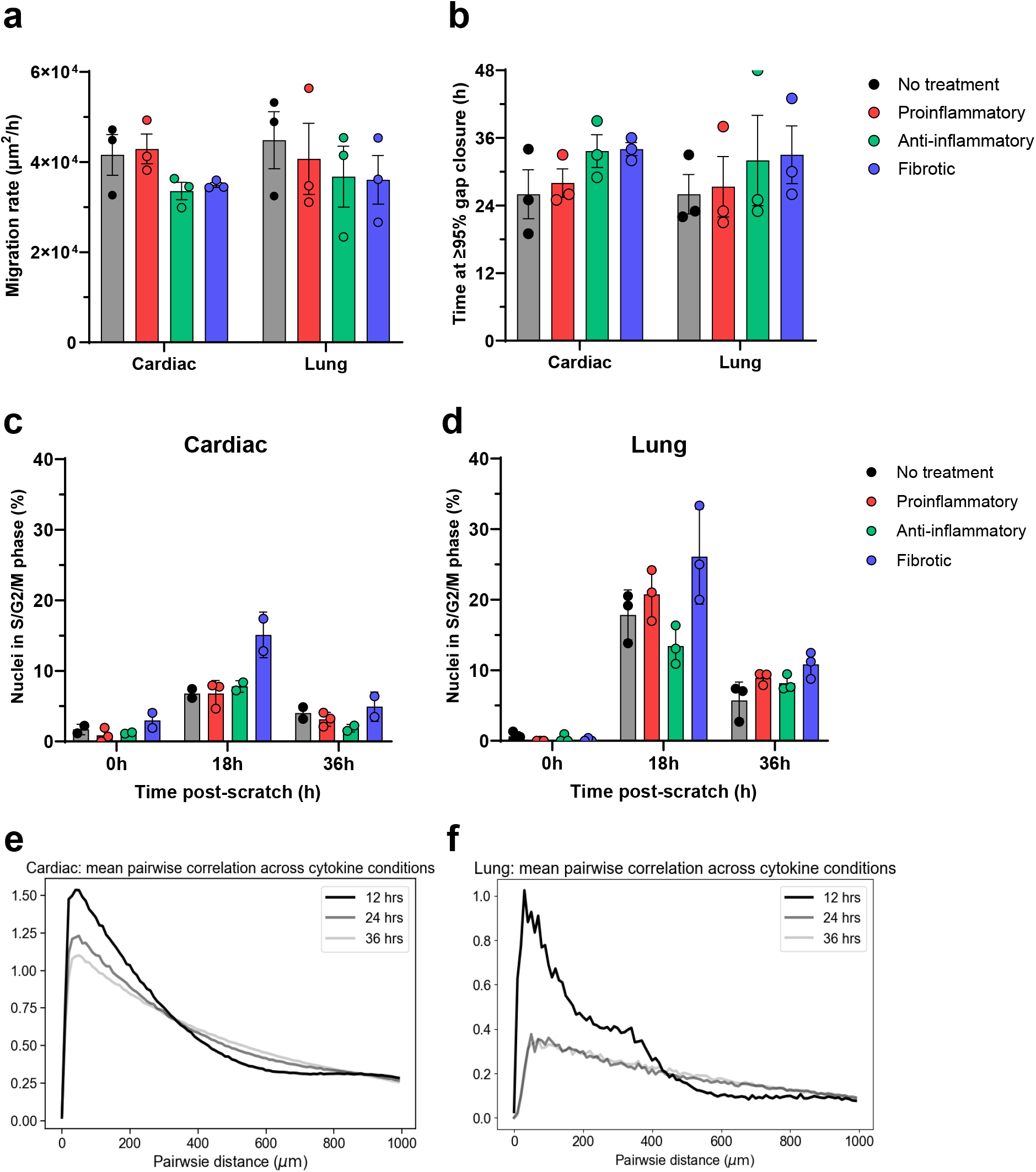
Fibroblast migration in an *in vitro* scratch assay. **(a)** Migration rate and **(b)** the time taken to achieve ≥95% gap closure by cardiac and lung fibroblasts in response to cytokine cocktails that broadly represent proinflammatory, anti-inflammatory and fibrotic microenvironments. Data is the mean of three biological replicates from independent experiments with SEM. No statistical significance was determined using 2-way ANOVA with Šídák’s post-hoc testing. **(c)** Percentage of cardiac and **(d)** lung Fucci2 fibroblasts detected in the S/G2/M cell cycle phase at 0-, 18-, and 36-hours post-scratch. Data is the mean of up to three technical replicates with SD. **(e)** Pairwise correlations of cardiac and **(f)** lung fibroblasts. Data is obtained from experimental results at 12-, 24-, and 36-hours post scratch, and averaged over all cytokine conditions.

**Figure S2.**
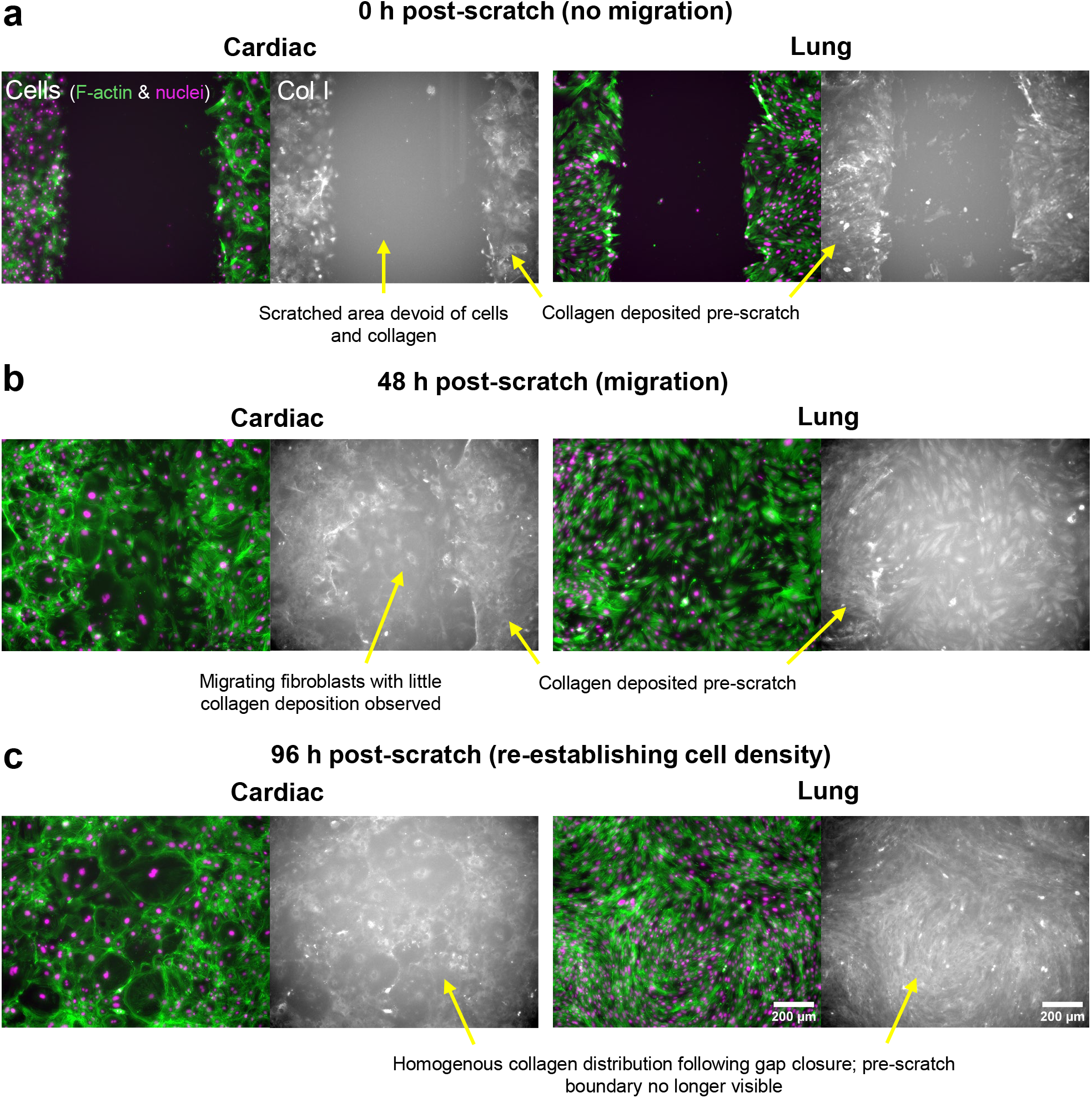
Collagen I staining during fibroblast migration in a scratch assay. Cardiac and lung fibroblasts were stained for collagen I at 0, 48, and 96 hours post-scratch to assess the timing of collagen I deposition relative to cell migration. Fibroblasts were stained and imaged directly on tissue-culture plastic multi-well plates and were not permeabilised prior to staining with an anti-collagen I antibody. **(a)** Prior to scratching, fibroblasts formed a confluent monolayer and deposited collagen I. The scratch removed both cells and collagen, creating a cell- and collagen-free wound bordered by collagen-rich regions. **(b)** During the first 48 hours post-scratch, fibroblasts migrated into the wound area. Although collagen synthesis was detectable at sites of migrating cells, no substantial collagen network formed within the wound area, leaving a clear boundary between the pre-scratch collagen and migrating cells. This suggests that fibroblasts prioritise migration over matrix deposition at this stage. **(c)** Following gap closure and re-establishment of high cell density in the wound, robust collagen deposition was observed, resulting in a uniform collagen distribution across the field of view and loss of the original pre-scratch boundary.

**Figure S3.**
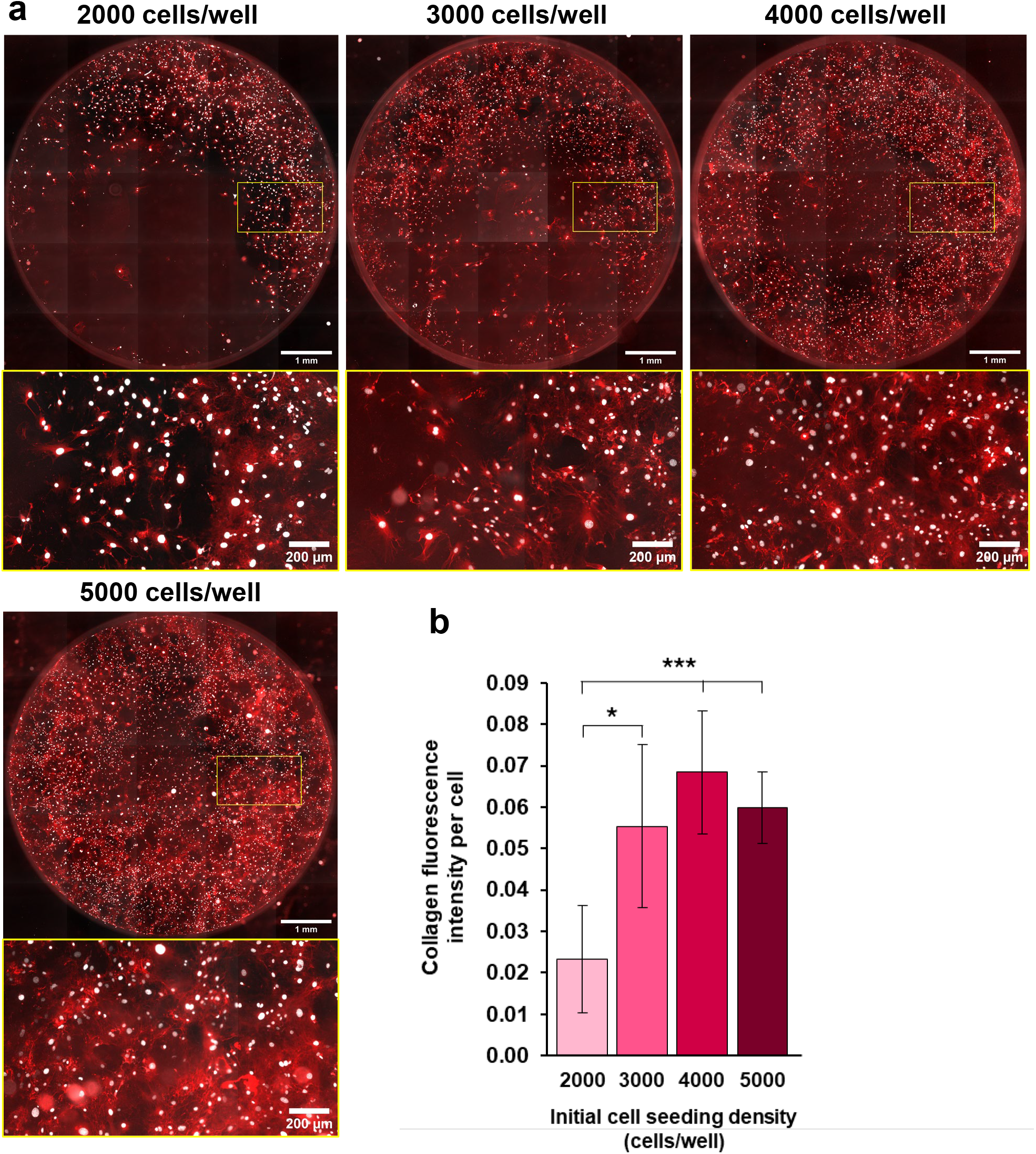
Collagen I deposition in response to different cell seeding densities. **(a)** Collagen I deposited by cardiac fibroblasts initially seeded at 2000, 3000, 4000, and 5000 cells/well in a 96-well plate and cultured in complete growth media with ascorbic acid (100 µg/mL) for 3 days. Cells were fixed and stained for collagen I followed by imaging of the entire well to display heterogenous cell and collagen distributions. **(b)** Collagen fluorescence intensity (measured by CTCF, corrected total collagen fluorescence) normalised to the number of cells per well to account for differences in cell density. Data is the mean of four replicates with standard deviation. Statistical significance was determined by a one-way ANOVA with Tukey’s post-hoc testing (* *p*<0.05, *** *p*<0.005).

**Figure S4.**
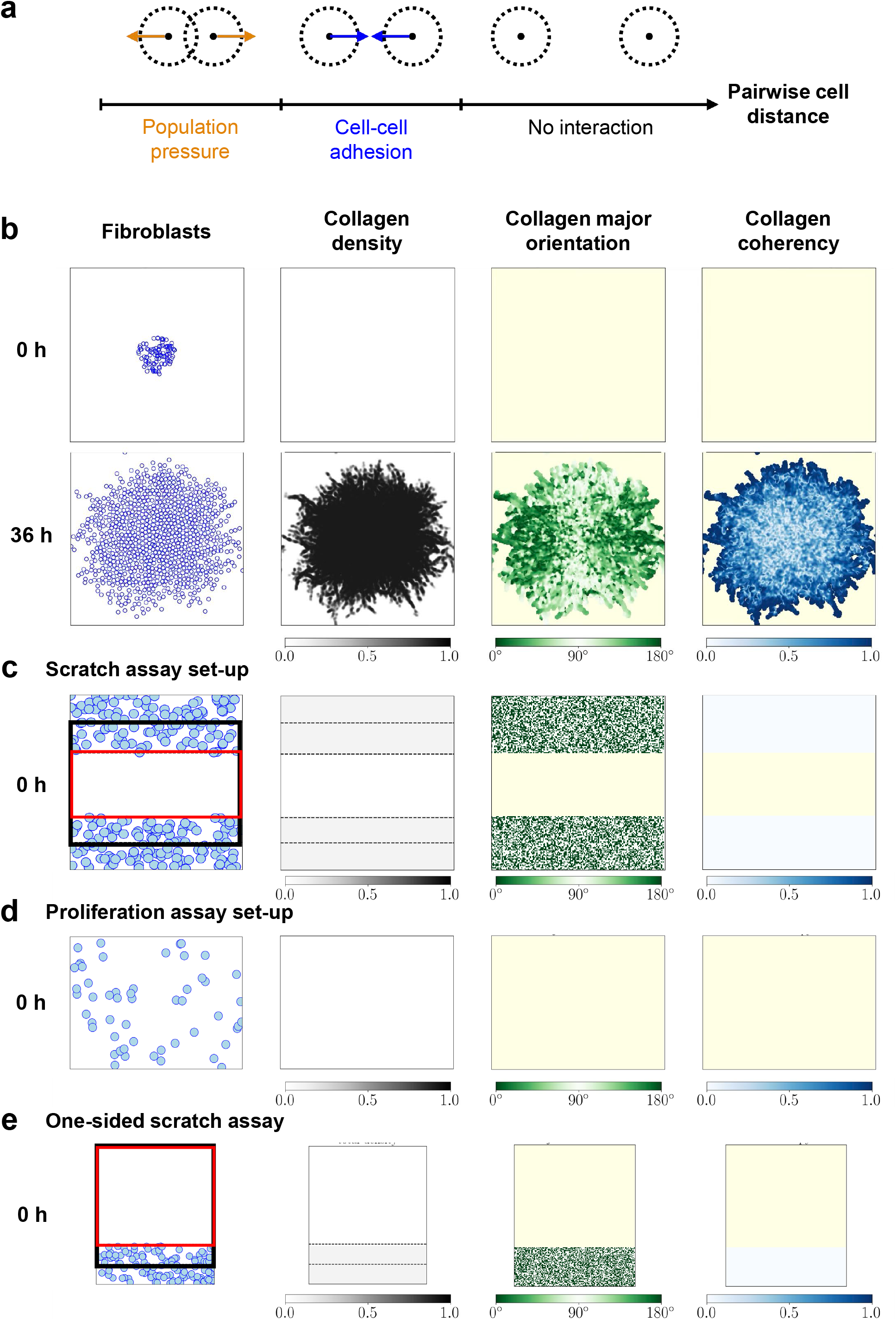
Schematic diagram of the computational model. **(a)** Pairwise cell-cell interactions are determined by the distance between fibroblasts. Fibroblasts are modelled as individual agents characterised by a central position (solid point) and an interaction length scale (dashed circle) defining their interaction range. Fibroblasts exert short-range repulsive forces (orange arrows) when in close proximity due to population pressure, adhesive forces (blue arrows) at intermediate distances, and exhibit no interaction when sufficiently far apart. **(b)** Simulated fibroblast and collagen distributions at 0 and 36 hours, under conditions of fixed random motility independent of collagen density and a constant collagen secretion rate regardless of cell density, as in Yin et al. [2025]. Fibroblasts are depicted as blue circles, with the circle radius representing the fibroblast radius. Regions devoid of collagen are coloured in pale yellow. Collagen properties are coloured as follows: total density in greyscale (darker indicates higher density), major orientation relative to the horizontal *x* axis in green (darker indicates more vertical alignment), and coherency in blue (darker indicates more aligned collagen). Initially, 100 fibroblasts are arranged in a densely packed, confluent circular disc at the centre of a [0, 540] × [0, 540] *μ*m^2^ domain with periodic boundary conditions, in the absence of any collagen. **(c)** Scratch assay initial simulation setup. Regions outside the wound area are initialised with collagen that is uniformly distributed in random directions with a total density of 0.1. The black box denoting FOV matches the experimental setup, while the red box encloses the wound region. We impose flux *y*-boundary conditions, so that the number of cells are fixed at confluency level throughout the simulation in the regions outside the FOV. We have periodic *x*-boundary conditions. **(d)** Proliferation assay initial simulation setup matching the experimental FOV. We impose periodic boundary conditions. **(e)** One-sided scratch assay initial simulation setup. Regions outside the wound area are initialised with collagen that is uniformly distributed in random directions with a total density of 0.1. The black box denoting FOV while the red box encloses the wound region. We impose flux *y*-boundary conditions, so that the number of cells are fixed at confluency level throughout the simulation in the regions outside the FOV. We have periodic *x*-boundary conditions.

**Figure S5.**
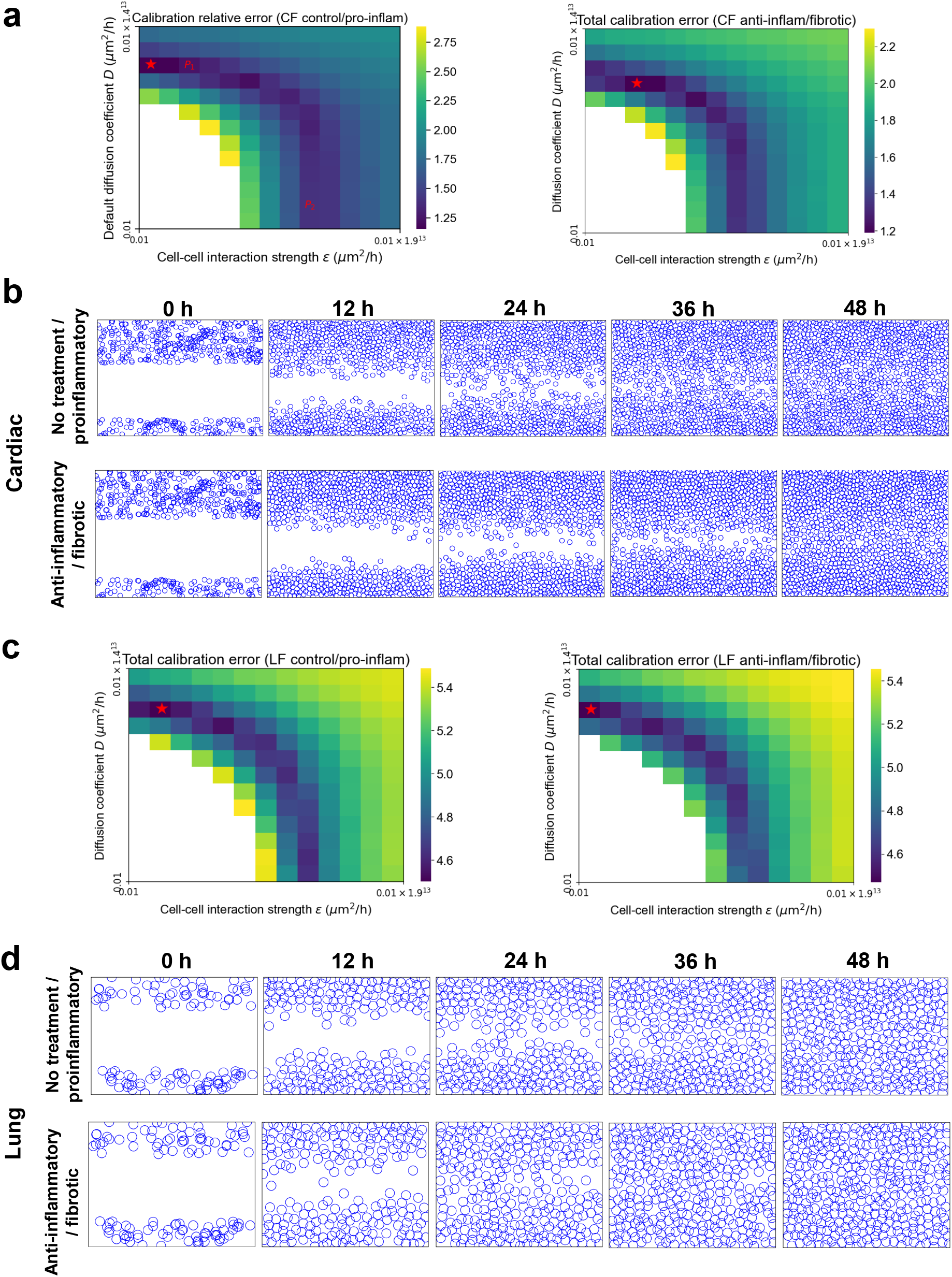
Optimal parameter identification for calibrating the model to experimental data. Total mean relative errors for **(a)** cardiac and **(c)** lung fibroblasts under control/proinflammatory and anti-inflammatory/fibrotic conditions. The red stars denote parameter values giving rise to the minimal relative errors. Snapshots of simulated migration dynamics for **(b)** cardiac and **(d)** lung fibroblasts using optimal parameter values for different cytokine conditions that best matched the experimental data.

**Figure S6.**
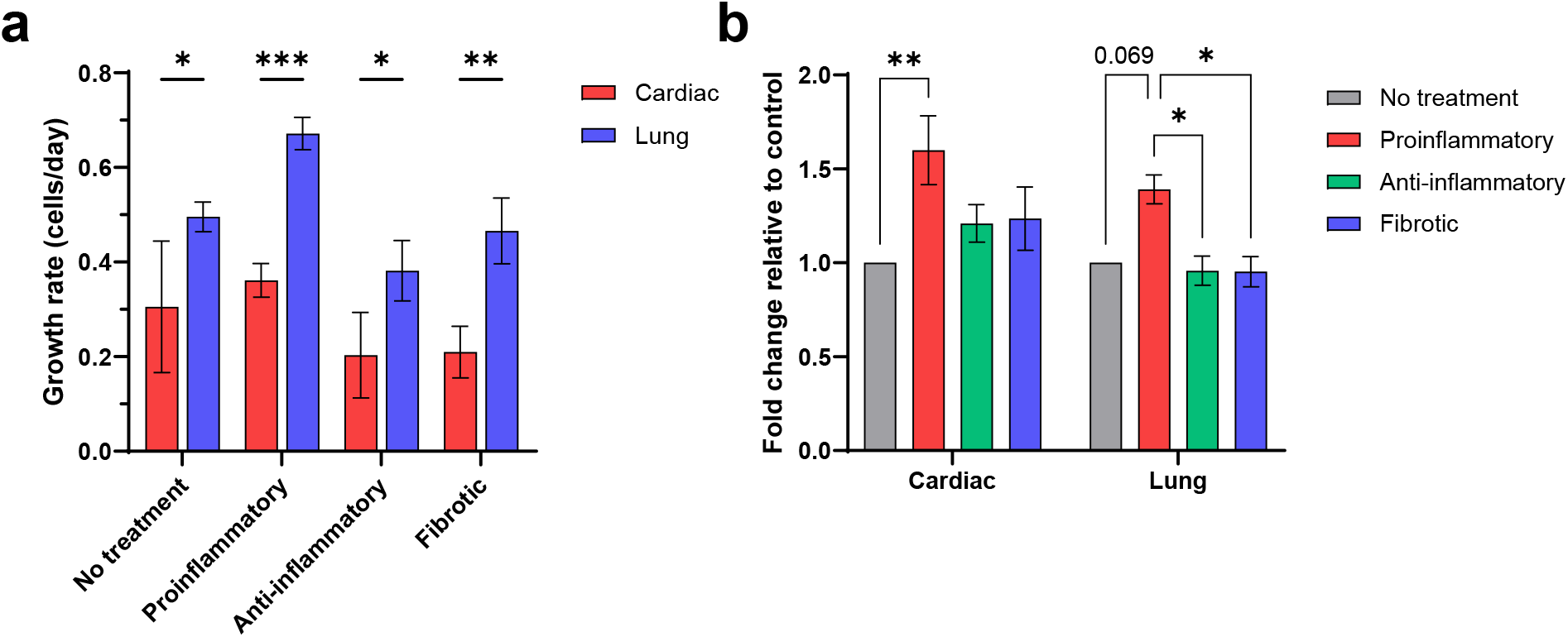
Growth rate and metabolic activity of cardiac and lung fibroblasts. **(a)** Growth rates of cardiac and lung fibroblasts cultured for 72 hours in different cytokine cocktails. **(b)** Corresponding cell metabolic activity in response to different cytokine cocktails. Raw measurement was the absorbance of WST-1 at 440 nm and data converted to fold change relative to control conditions. Data is the mean of (a) three and (b) four biological replicates with SEM. Statistical significance was determined by ANOVA with Tukey’s post-hoc testing (* *p*<0.05, ** *p*<0.01, *** *p*<0.005).

**Figure S7.**
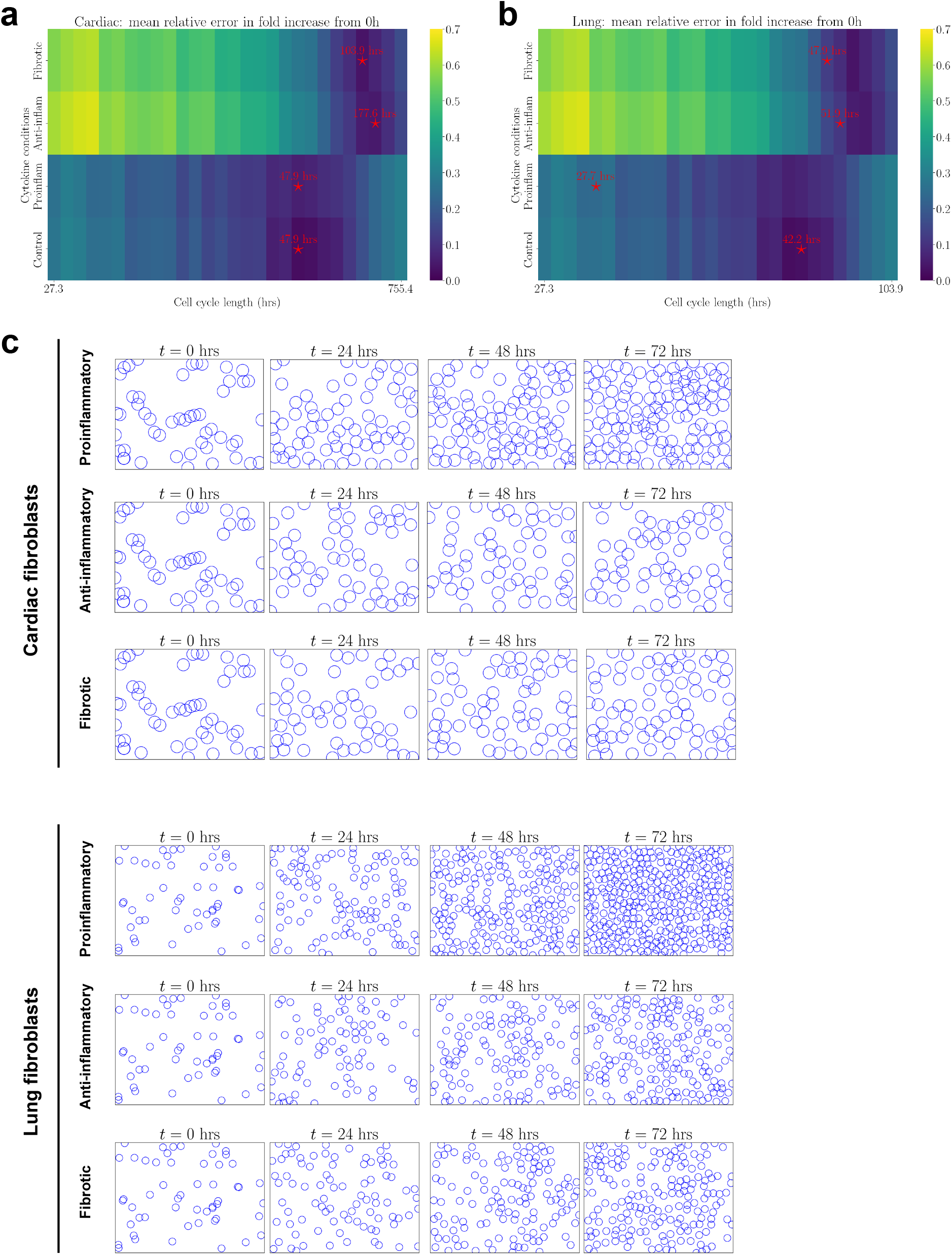
Calibration the computational model to the proliferation assay data. Mean relative errors in fold increase from the 0 hour for **(a)** cardiac and **(b)** lung fibroblasts in different cytokine conditions, respectively. The cell cycle lengths corresponding to the minimum relative errors are labelled using red stars, and corresponding dynamics are shown in **(c)**.

**Figure S8.**
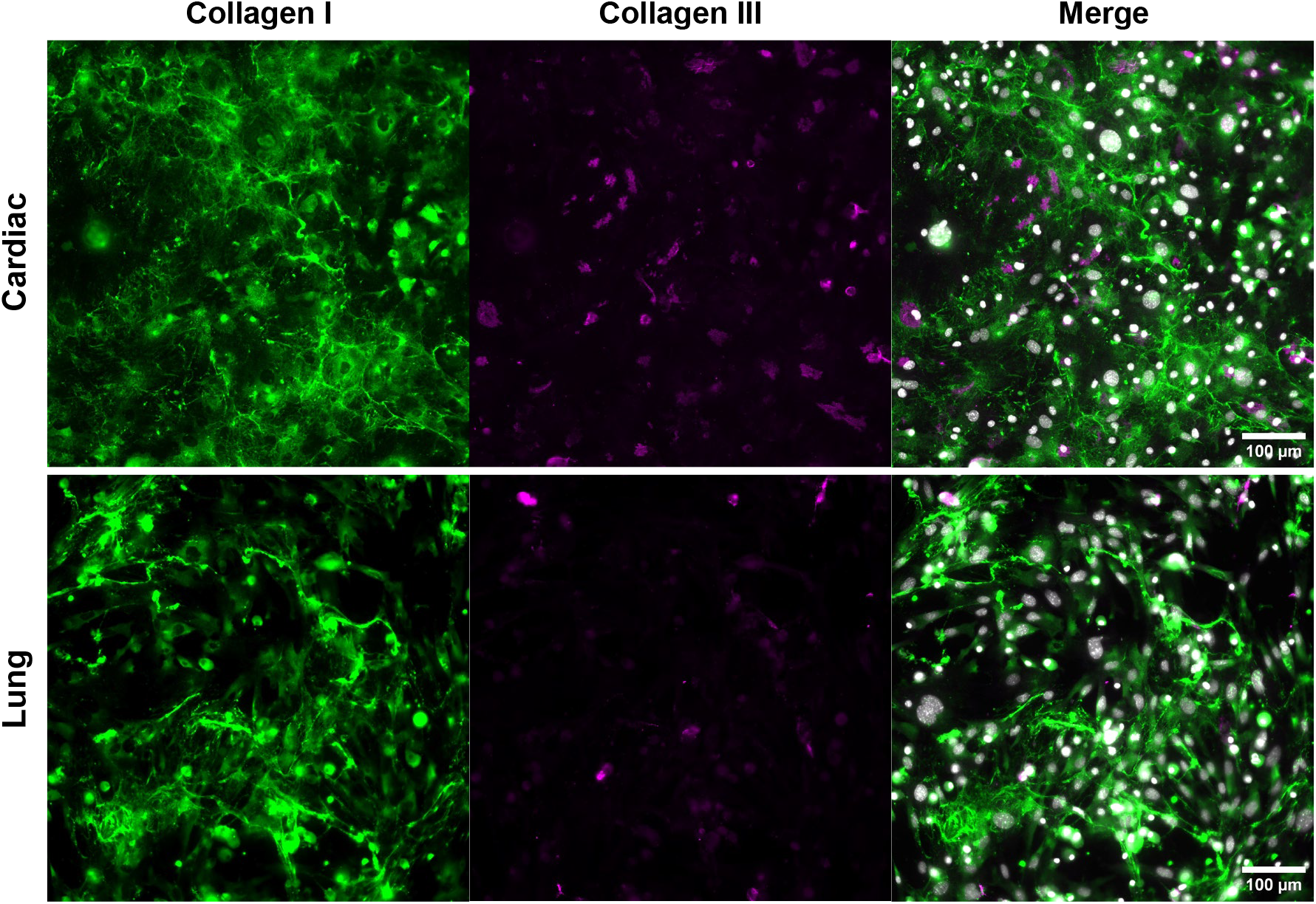
Collagen production by cardiac and lung fibroblasts. Collagen I and III deposited by cardiac and lung fibroblasts after 72 hours. Fibroblasts were cultured in media with ascorbic acid (100 µg/mL).

**Figure S9.**
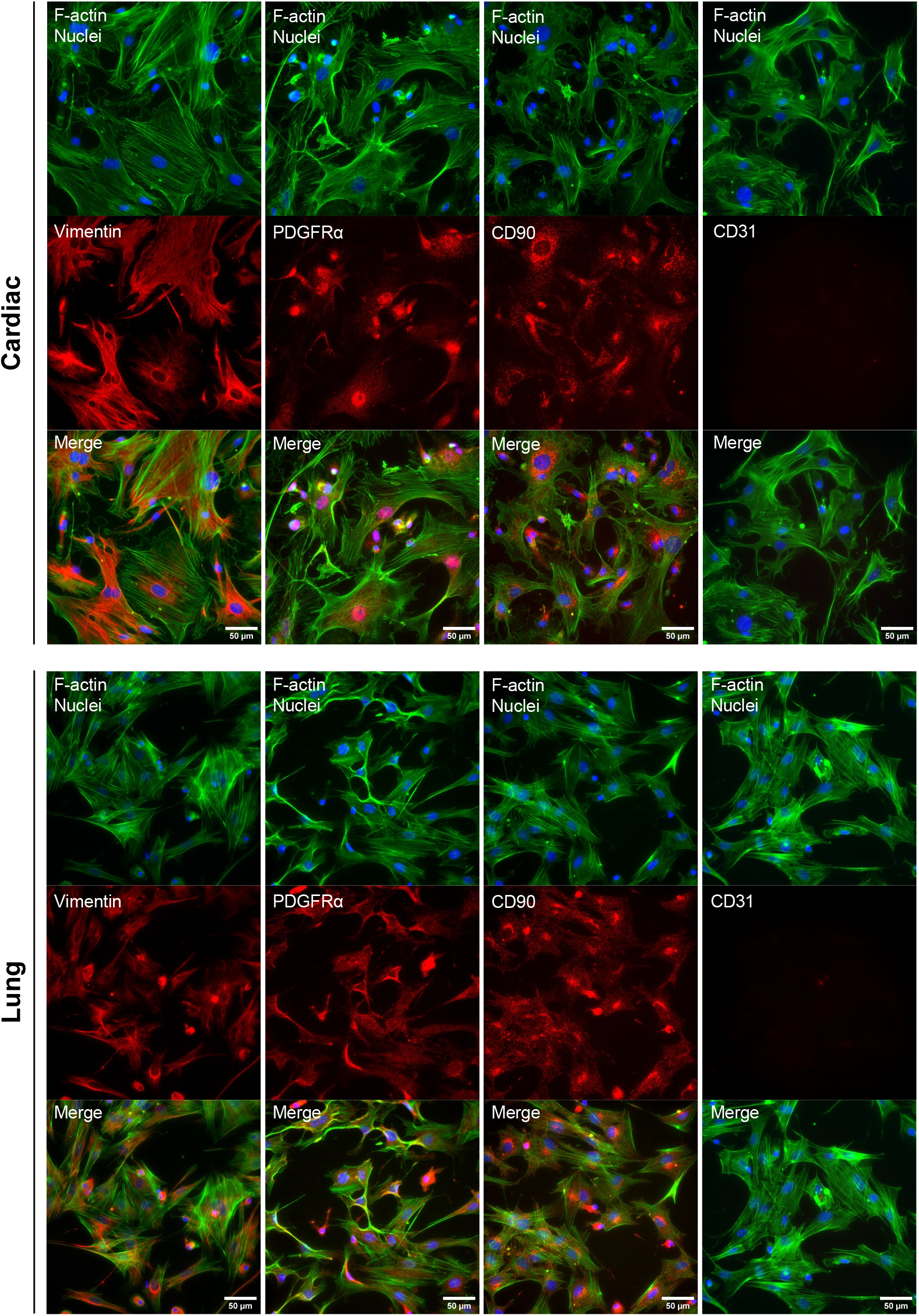
Isolated primary murine cells express markers associated with a fibroblast phenotype. Primary cells isolated from cardiac and lung tissue expressed vimentin, PDGFRα and CD90 and did not express the endothelial marker CD31.

## Footnotes

1 We set ***X***^*i*^(*t*-*m*) = ***X***^*i*^ (0), the initial position for cell *i* = 1, …, *N*, when *t* < *m*.

## Notes

https://github.com/YuanYIN99/OrganSpecificFibrosis.git

### Competing Interest Statement

The authors have declared no competing interest.

